# Targeting oncogenic K-Ras mutants with a small-molecule degrader through Nedd4-1

**DOI:** 10.1101/2024.04.26.591418

**Authors:** Taoling Zeng, Tingting Jiang, Baoding Zhang, Ting Zhang, Wanjun Dai, Xun Yin, Yunzhan Li, Caiming Wu, Yaying Wu, Ximin Chi, Xianming Deng, Hong-Rui Wang

## Abstract

K-Ras mutations represent a most prevalent oncogenic alteration in human cancers. Despite of tremendous efforts, it remains a big challenge to develop inhibitors that can target the oncogenic K-Ras mutants, especially mutants without specific active or charged side chains such as K-Ras^G12V^. Here, taking advantage of our previous finding that Nedd4-1 is a bona fide E3 ubiquitin ligase for wild-type Ras proteins, we developed a compound XMU-MP-9 that can promote ubiquitination and degradation of various K-Ras mutants including K-Ras^G12V^, and significantly inhibit proliferation and tumor development of K-Ras mutant harboring cells. Mechanistically, XMU-MP-9 acts as a bifunctional compound to bind the C2 domain of Nedd4-1 and an allosteric site of K-Ras to enhance Nedd4-1 and K-Ras interaction, and to induce a conformational change of Nedd4-1/K-Ras complex to allow Nedd4-1 targeting K128 of K-Ras for ubiquitination. Hence, our study presents a robust strategy to develop small-molecule degrader of K-Ras mutants, and also sheds light on the development of small-molecule degraders for H-Ras and N-Ras mutants.

## Introduction

Oncogenic mutations of Ras are associated with about 30% of all human cancers^1^. Mutations in Ras predominantly occur at G12, G13, and Q61, resulting in a persistence of the guanosine triphosphate (GTP)-bound state of Ras by impairing the intrinsic and GTPase-activating proteins (GAPs)-stimulated GTP hydrolysis reaction^2–4^. Oncogenic Ras mutants can trigger constitutive activation of downstream signaling pathways such as Raf/MEK/ERK signal transduction cascade, leading to uncontrolled cell proliferation, transformation, differentiation, and enhanced cell survival^5,6^. Although mutations of different Ras isoforms (K-Ras, H-Ras, and N-Ras) are associated with different types of tumors in a non-random distribution pattern, K-Ras is mutated most frequently among the three human Ras proteins^7^. Indeed, targeting K-Ras mutants for cancer therapy has garnered most attention from the researchers.

Despite of decades of extensive endeavors, targeting Ras mutants for cancer therapy remains intractable^1,8–10^. Because Ras proteins bind GTP with picomolar affinity, early approach on developing competitive antagonists of GTP binding achieved no success^1,10^. Meanwhile, the challenge of directly targeting Ras is also raised by no deep hydrophobic pockets on the surface of Ras for small-molecule compound binding^11^, which led to a perception that Ras proteins are undruggable^1,12^. However, given the prevalence of Ras mutations in cancers, targeting mutated Ras proteins is still drawing very much attention^1,8–10,12,13^. Recent studies reported designs of pan-Ras inhibitors that can simultaneously bind to two or more adjacent sites on Ras proteins, therefore prevent effector protein binding^14,15^. Kim et al. developed a pan-K-Ras inhibitor that can block nucleotide exchange of wild-type K-Ras and a broad range of K-Ras mutants^16^. Another exciting approach was to selectively target Ras^G12C^ mutant by covalently linking an inhibitor to the Cys12^17–20^. Hallin et al. developed a non-covalent K-Ras^G12D^-selective inhibitor taking advantage of the acidic side chain of Asp12, obtaining a 700-fold selectivity for binding to K-Ras^G12D^ vs. wild-type K-Ras^21^. Recent studies also developed tri-complex Ras inhibitors, which may induce tri-complex formation of immunophilin protein cyclophilin A, compound, and Ras, therefore blocks interactions of downstream effectors with Ras^22–25^. However, targeting the active state of K-Ras mutants lacking specific active or charged side groups like K-Ras^G12V^ with small-molecule drugs still poses challenges.

Recent advances in the development of targeted protein degradation via ubiquitination machinery provide an exciting strategy for targeting undruggable proteins^26,27^. The proximity-inducing agents that can induce a close proximity between an E3 ubiquitin ligase and its non-native target protein include molecular glues and bifunctional compounds^28^. Proteolysis targeting chimeras (PROTACs) are the most advanced type of bifunctional small-molecule degraders, composed by a ligand for binding of protein of interest, another ligand specific for an E3 ubiquitin ligase, and a linker to connect the two ligands^29^. Our previous study showed that Nedd4-1 acts as a general E3 ubiquitin ligase for all three major forms of Ras (K-Ras, H-Ras, and N-Ras) to control their abundance in normal cells^30^. Nedd4-1 preferably targets guanosine diphosphate (GDP)-bound Ras but not GTP-bound form for degradation because binding of GTP causes a conformational change of ubiquitination site K5 to evade Nedd4-1-mediated ubiquitination. Therefore, oncogenic Ras mutants that constitutively bind GTP are resistant to the ubiquitination mediated by Nedd4-1. Interestingly, even though Nedd4-1 does not promote ubiquitination of oncogenic Ras mutants, it binds these Ras mutants with a similar affinity to that of wild-type Ras^30^. In this study, taking advantage of the pre-existing interaction of K-Ras mutants and Nedd4-1, we successfully identified a small-molecule compound XMU-MP-9 that can act as a bifunctional agent to facilitate Nedd4-1-mediated ubiquitination and degradation of various oncogenic K-Ras mutants, thereby inhibits the proliferation and tumor development of K-Ras mutant harboring cancer cells.

## Results

### XMU-MP-9 promotes Nedd4-1-mediated K-Ras^G12V^ degradation

Our previous study demonstrated that Nedd4-1 is a bona fide E3 ubiquitin ligase for wild-type Ras proteins. Although oncogenic Ras mutants are resistant to Nedd4-1-mediated ubiquitination, they still can bind to Nedd4-1. We speculated that it might be possible to enable Nedd4-1 to ubiquitinate oncogenic Ras mutants using a small-molecule compound. This compound is expected to promote degradation of oncogenic Ras mutants in the presence of Nedd4-1, thereby potentially inhibiting the proliferation of wild-type cells but not Nedd4-1 knockout cells. Therefore, in attempting to identify small-molecule compound(s) that can induce Nedd4-1-mediated degradation of oncogenic K-Ras mutants, we utilized SW620, a human colorectal cancer cell line that harbors K-Ras^G12V^, and performed a high-throughput screen to look for compounds having more growth inhibitory potency to wild-type (Nedd4-1^+/+^) than to Nedd4-1 knockout (Nedd4-1^-/-^) SW620 cells. For this screen, cells seeded in 96-well plates were treated with chemical compounds for 4 days and then subjected to cell viability examination. Compounds with at least 20% more inhibitory effect on Nedd4-1^+/+^ cells than that on Nedd4-1^-/-^ cells were selected for further characterization (Fig. 1a). We used an in-house compound library containing a total of 4,480 compounds and divided them into 1,120 groups by mixing 4 different compounds into each group. In the first-round screening, 17 groups (68 compounds in total) met our selection criteria. The 68 compounds were then applied for a second-round screen and 13 compounds were identified with a preference to inhibit proliferation of wild-type SW620 cells (Fig. 1b, c). Among them, one compound showed a specific effect on inducing decrease of endogenous K-Ras^G12V^ protein levels in Nedd4-1^+/+^ but not Nedd4-1^-/-^ cells (Extended Data Fig. 1a), and was named XMU-MP-9 (Fig. 1d) in our following studies. XMU-MP-9 could induce decrease of endogenous K-Ras^G12V^ protein levels in SW620 cells as early as 12 hours treatment and reached a maximum effect at 36 to 48 hours (Extended Data Fig. 1b). XMU-MP-9 not only presented a more potent effect on inhibiting growth of Nedd4-1^+/+^ SW620 cells in 2-D culture and 3-D soft agar in comparison to Nedd4-1^-/-^ SW620 cells (Fig. 1e, f), but also showed a Nedd4-1-dependent decrease of endogenous K-Ras^G12V^ protein levels in a dose-dependent manner (Fig. 1g and Extended Data Fig. 1c).

**Figure 1.**
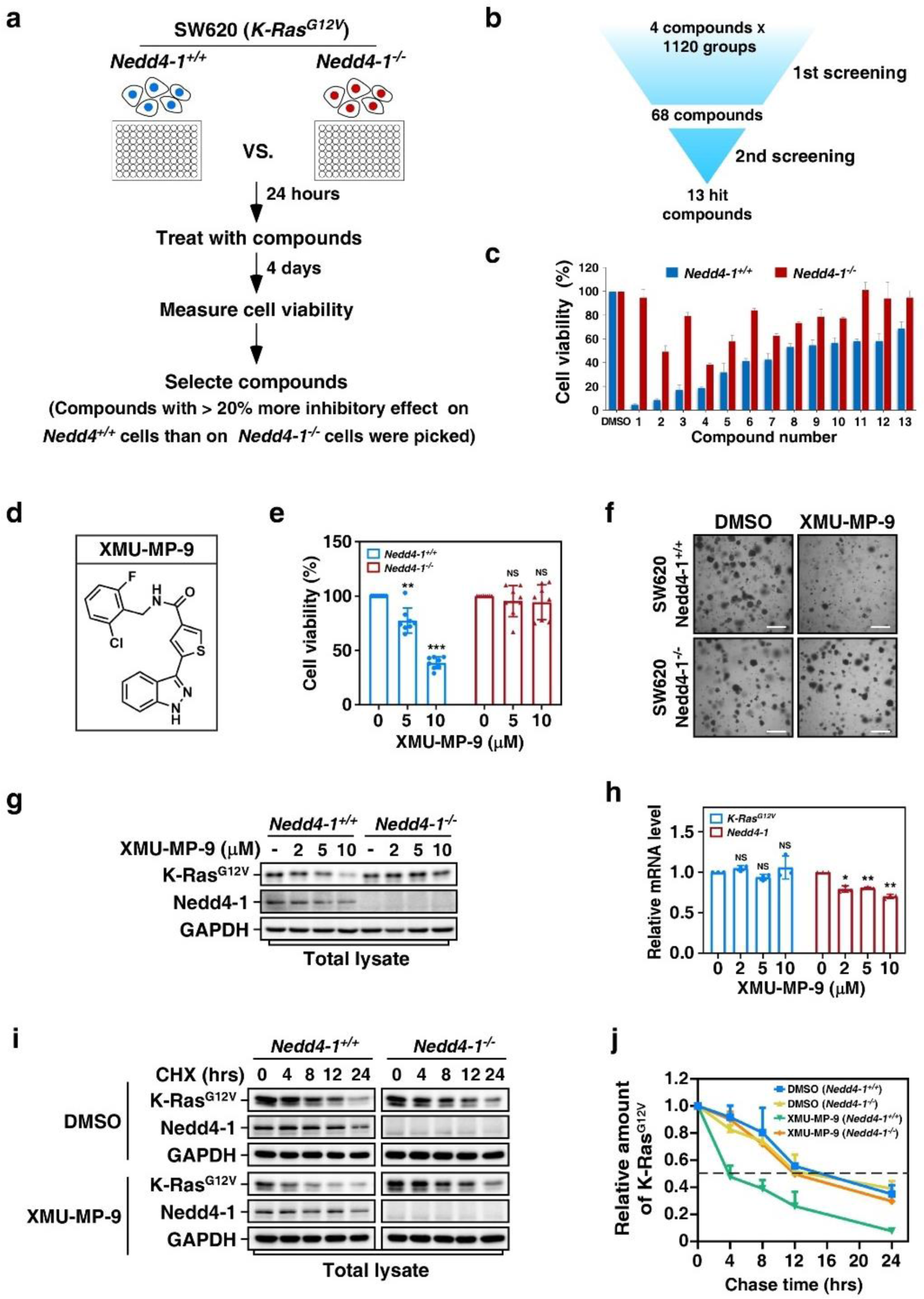
XMU-MP-9 induces Nedd4-1-dependent K-Ras^G12V^ degradation. **a,** Schematic of high-throughput screening for compounds preferably inhibiting growth of wild-type (Nedd4-1^+/+^) but not Nedd4-1 knockout (Nedd4-1^-/-^) SW620 cells. **b,** Screening of the compound library for the hits. **c,** Identification of individual compounds with preference for inhibiting growth of wild-type SW620 cells. Nedd4-1^+/+^ and Nedd4-1^-/-^ SW620 cells seeded in 96-well plates were treated 4 days with 10 μM of each compound selected from the screen and then subjected to cell viability assay. Data were presented as mean ± SD of three individual experiments. **d,** Chemical structure of XMU-MP-9. **e, f,** XMU-MP-9 specifically inhibits growth of wild-type but not Nedd4-1 knockout SW620 cells. Nedd4-1^+/+^ and Nedd4-1^-/-^ SW620 cells were seeded in 2-D culture and treated with indicated concentrations of XMU-MP-9 for 4 days before subjected to cell viability assay (**e**), or seeded in 3-D soft agar and treated 10 μM XMU-MP-9 or DMSO as control for 2 weeks (**f**). Cell viabilities were determined and plotted as mean ±SD (n = 8). *P* values were determined by one-way ANOVA followed by Tukey test. * *p* < 0.05; ** *p* < 0.01; *** *p* < 0.001; NS, not significant. Scale bar indicates 100 μm. **g,** XMU-MP-9 induces decrease of steady-states levels of K-Ras^G12V^ in Nedd4-1^+/+^ but not Nedd4-1^-/-^ SW620 cells. Nedd4-1^+/+^ and Nedd4-1^-/-^ SW620 cells treated 48 h with XMU-MP-9 as indicated were subjected to immunoblotting assay. **h,** Effects of XMU-MP-9 on transcription of K-Ras^G12V^ and Nedd4-1. SW620 cells treated 48 h with XMU-MP-9 as indicated were subjected to qRT-PCR assay to determine the mRNA levels of K-Ras^G12V^ and Nedd4-1. The results are presented as mean ±SD of 3 independent experiments. *P* values were determined by one-way ANOVA followed by Tukey test. * *p* < 0.05; ** *p* < 0.01; *** *p* < 0.001; NS, not significant. **i, j,** XMU-MP-9 promotes Nedd4-1-dependent degradation of K-Ras^G12V^. Nedd4-1^+/+^ and Nedd4-1^-/-^ SW620 cells treated with 10 μM XMU-MP-9 and 100 μg/ml cycloheximide (CHX) were collected at indicated time points and subjected to immunoblotting assay to determine the protein levels (**i**). Relative amounts of K-Ras^G12V^ at each time point to the level at time 0 were plotted. The results are presented as mean ±SD of 3 independent experiments (**j**).

We confirmed that the decrease of K-Ras^G12V^ was not through change of transcription as treatment of XMU-MP-9 did not affect the mRNA levels of K-Ras^G12V^ (Fig. 1h). It is noteworthy that treatment of XMU-MP-9 induced decrease of mRNA levels of Nedd4-1 (Fig. 1h), which agrees with our previous report that Ras signaling is required for Nedd4-1 transcription^30^. Co-treatment with lysosome inhibitor chloroquine but not proteasome inhibitor MG-132 significantly blocked the decrease of endogenous K-Ras^G12V^ induced by XMU-MP-9 (Extended Data Fig. 1d), indicating a lysosomal degradation of K-Ras^G12V^. Moreover, in agreement with our previous study that K-Ras oncogenic mutants are resistant to Nedd4-1-mediated degradation, knockout of Nedd4-1 in SW620 cells did not significantly affect protein half-life of K-Ras^G12V^; however, treatment of XMU-MP-9 significantly decreased the K-Ras^G12V^ protein half-life only in the presence of Nedd4-1 (Fig. 1i, j), indicating that XMU-MP-9-promoted K-Ras^G12V^ degradation is indeed dependent on Nedd4-1.

### XMU-MP-9 promotes Nedd4-1-mediated ubiquitination of K-Ras^G12V^ on K128

We next conducted ubiquitination assays to verify the effect of XMU-MP-9 on promoting ubiquitination of K-Ras^G12V^. Treatment of XMU-MP-9 significantly enhanced ubiquitination of endogenous K-Ras^G12V^ in wild-type SW620 cells in a dose dependent manner, whereas knockout of Nedd4-1 totally blocked the XMU-MP-9-promoted K-Ras^G12V^ ubiquitination (Fig. 2a). To avoid potential contamination of K-Ras binding proteins, we performed 2× anti-Flag immunoprecipitation (IP) to confirm the effect of XMU-MP-9 on promoting ubiquitination of K-Ras^G12V^. Exogenously expressed Flag-tagged K-Ras^G12V^ was subjected to anti-Flag immunoprecipitation, eluted by boiling in 1% SDS, then diluted in TNTE lysis buffer and re-immunoprecipitated with anti-Flag IP (2×IP) (Fig. 2b). In addition, we performed an *in vitro* ubiquitination assay using bacterially expressed Nedd4-1 and Flag-tagged K-Ras^G12V^, in which XMU-MP-9 drastically enhanced the Nedd4-1-mediated K-Ras^G12V^ ubiquitination (Fig. 2c), demonstrating a clear evidence of XMU-MP-9 in directly assisting Nedd4-1-mediated ubiquitination of K-Ras^G12V^. Notably, auto-ubiquitination of Nedd4-1 was not enhanced by treatment of XMU-MP-9 (Fig. 2d), suggesting that XMU-MP-9 does not function as an activator to increase the catalytic activity of Nedd4-1.

**Figure 2.**
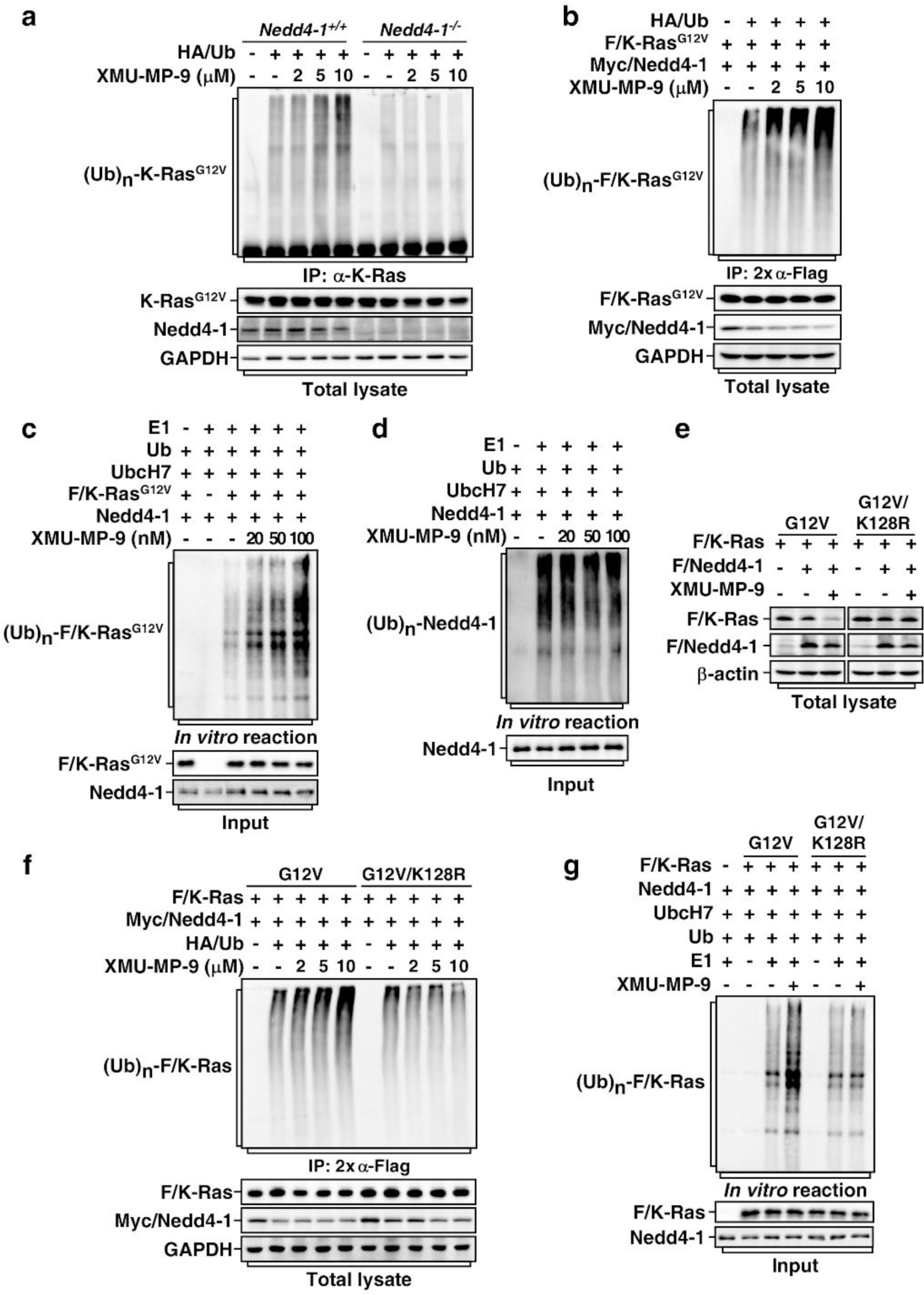
XMU-MP-9 promotes Nedd4-1-mediated ubiquitination of K-Ras^G12V^. **a**, XMU-MP-9 promotes ubiquitination of endogenous K-Ras^G12V^ in wild-type but not Nedd4-1 knockout cells. Nedd4-1^+/+^ and Nedd4-1^-/-^ SW620 cells transduced with lentivirus encoding HA-tagged ubiquitin (HA/Ub) were pre-treated 1 h with 120 μM chloroquine and then 1 h with indicated concentrations of XMU-MP-9 before subjected to anti-K-Ras immunoprecipitation (IP). Ubiquitin-conjugated K-Ras^G12V^ were detected by immunoblotting assay. **b,** XMU-MP-9 promotes Nedd4-1-mediated ubiquitination of exogenously expressed K-Ras^G12V^. HEK293T cells transfected with indicated combinations of HA/Ub, Flag-tagged K-Ras^G12V^ (F/K-Ras^G12V^), and Myc-tagged Nedd4-1 (Myc/Nedd4-1) were treated same as in (**a**) and then subjected to 2× anti-Flag IP. Ubiquitin-conjugated F/K-Ras^G12V^ was detected by immunoblotting with anti-HA antibody. **c,** XMU-MP-9 promotes Nedd4-1-mediated ubiquitination of K-Ras^G12V^ *in vitro*. Nedd4-1 and F/K-Ras^G12V^ purified from bacteria were subjected to an *in vitro* ubiquitination assay in the presence or absence of indicated concentrations of XMU-MP-9. Reactions were stopped by boiling in 1% SDS and ubiquitination products were subjected to immunoprecipitation with anti-Flag antibodies. Ubiquitinated F/K-Ras^G12V^ was detected by immunoblotting assay with anti-Ub antibody. **d**, XMU-MP-9 does not enhance auto-ubiquitination of Nedd4-1. Nedd4-1 purified from bacteria was subjected to an *in vitro* ubiquitination assay with or without XMU-MP-9 as indicated. Ubiquitinated Nedd4-1 was detected by immunoblotting assay with anti-Ub antibody. **e**, K128R mutation blocks XMU-MP-9-induced degradation of F/K-Ras^G12V^. HEK293T cells transfected with Flag-tagged Nedd4-1 (F/Nedd4-1) and F/K-Ras^G12V^ or F/K-Ras^G12V/K128R^ as indicated were treated with or without XMU-MP-9 (10 μM, 48 h) and then subjected to immunoblotting assay. **f,** K128R mutation blocks XMU-MP-9-induced ubiquitination of F/K-Ras^G12V^ in cells. HEK293T cells transfected with indicated combinations of HA/Ub, Myc/Nedd4-1, F/K-Ras^G12V^, and F/K-Ras^G12V/K128R^ were treated same as in (**b**). **g,** K128R mutation blocks Nedd4-1-mediated ubiquitination of F/K-Ras^G12V^ *in vitro*. Nedd4-1, F/K-Ras^G12V^, and F/K-Ras^G12V/K128R^ purified from bacteria were subjected to an *in vitro* ubiquitination assay as in (**c**).

To identify the ubiquitination site(s) of K-Ras^G12V^, we analyzed ubiquitinated K-Ras^G12V^ obtained from the *in vitro* ubiquitination assay in the presence of Nedd4-1 and XMU-MP-9 by mass spectrometry. Only K128 of K-Ras^G12V^ was detected with ubiquitin conjugation (Extended Data Fig. 2a). To confirm this, we examined all the Lys-to-Arg mutations of K-Ras^G12V^ and found that only K128R mutation totally abolished the XMU-MP-9-promoted K-Ras^G12V^ degradation (Fig. 2e and Extended Data Fig. 2b). Accordingly, K128R mutation substantially blocked the Nedd4-1-mediated ubiquitination of K-Ras^G12V^ promoted by XMU-MP-9 both in cells and *in vitro* (Fig. 2f, g), indicating that XMU-MP-9 primarily assists Nedd4-1 in attaching ubiquitin to K128 of K-Ras^G12V^.

### XMU-MP-9 functions as a bifunctional compound for Nedd4-1 and K-Ras^G12V^

Next, we sought to explore the mechanism of XMU-MP-9 in assisting the ubiquitination of K-Ras^G12V^ mediated by Nedd4-1. Because XMU-MP-9 directly promotes Nedd4-1-mediated K-Ras^G12V^ ubiquitination and degradation without increasing the catalytic activity of Nedd4-1, we speculated that it might act as a small-molecule degrader to help Nedd4-1 to ubiquitinate K-Ras^G12V^. We therefore examined the bindings of XMU-MP-9 to Nedd4-1 and K-Ras^G12V^ using microscale thermophoresis (MST)^31^. Interestingly, XMU-MP-9 interacted with both K-Ras^G12V^ and Nedd4-1 with similar binding affinities (*K*_d_ = 18.4 nM and *K*_d_ = 25.4 nM, respectively) (Fig. 3a, b), and it could dose-dependently increase the binding affinity between K-Ras^G12V^ and Nedd4-1 (Fig. 3c). We further carried out *in vitro* nickel (Ni) affinity pull-down experiment and found that XMU-MP-9 dose-dependently enhanced the interaction between His-K-Ras^G12V^ and Nedd4-1 (Fig. 3d). Hence, XMU-MP-9 functions as a bifunctional compound to bind both Nedd4-1 and K-Ras^G12V^, and might also induce a conformational change of the Nedd4-1/K-Ras^G12V^ complex to allow Nedd4-1 to target K128 of K-Ras^G12V^ for ubiquitination (Fig. 2f, g).

**Figure 3.**
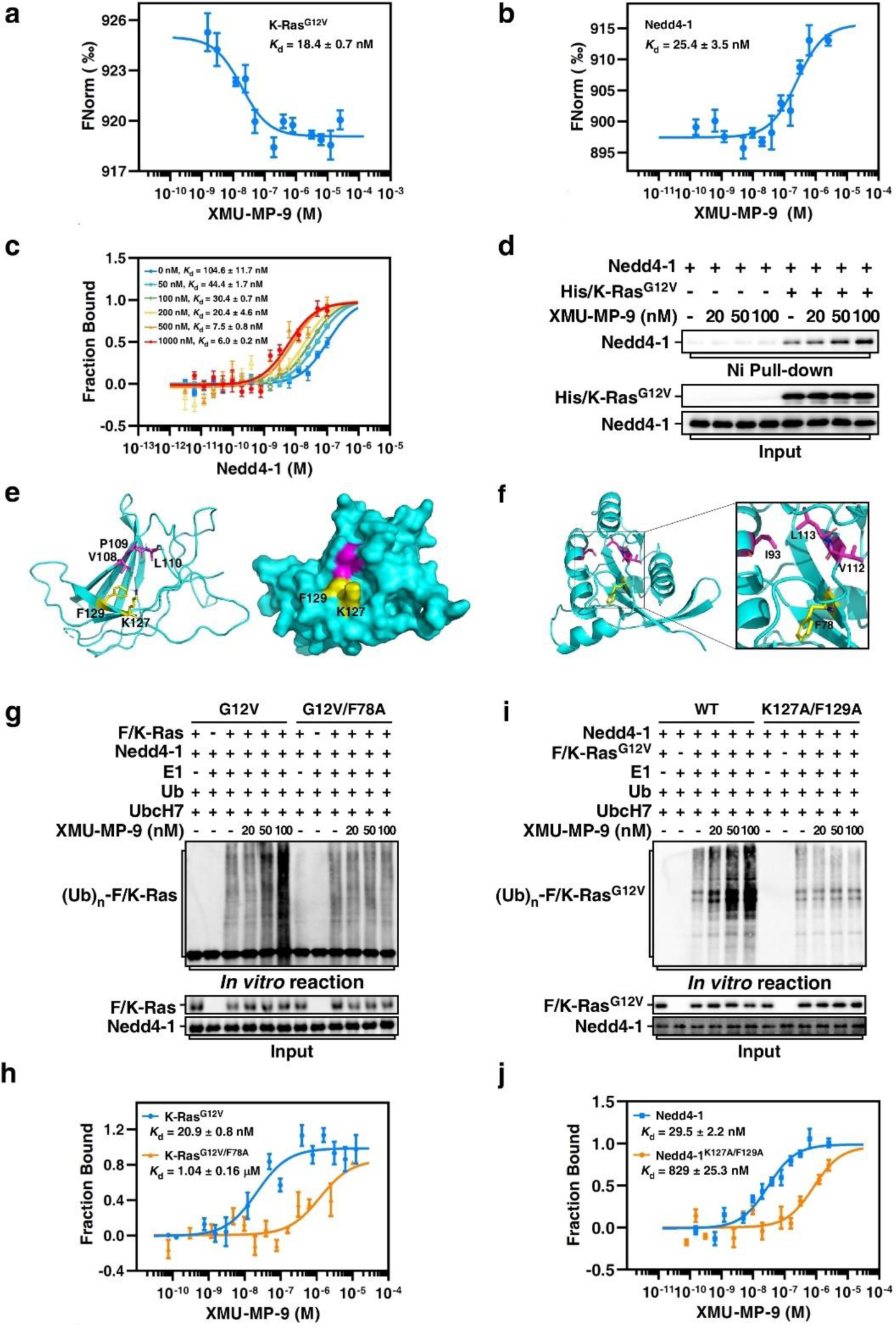
XMU-MP-9 acts as a bifunctional compound for Nedd4-1 and K-Ras^G12V^. **a,** XMU-MP-9 binds to K-Ras^G12V^. Red-NHS-labeled K-Ras^G12V^ (10 nM) with indicated concentrations of XMU-MP-9 were applied for microscale thermophoresis (MST) assays. Data of normalized fluorescence (FNorm in [‰]) were presented as mean ± SD of three individual experiments. The dissociation curve was fitted to the data to calculate the *K*_d_ value. **b,** XMU-MP-9 binds to Nedd4-1. Same as in (**a**) except Red-NHS-labeled Nedd4-1 instead of K-Ras^G12V^ was used. **c,** Presence of XMU-MP-9 increases binding affinity of Nedd4-1 to K-Ras^G12V^. Red-NHS-labeled K-Ras^G12V^ (10 nM) was mixed with indicated concentrations of Nedd4-1 in the presence of different concentrations of XMU-MP-9 as indicated for MST assays. Data of fraction bound were presented as mean ±SD of three individual experiments. **d,** XMU-MP-9 enhances interaction between K-Ras^G12V^ and Nedd4-1. Bacterially expressed and purified His-tagged (His/K-Ras^G12V^) and Nedd4-1 were subjected to nickel (Ni) affinity pull-down assay. Associated Nedd4-1 was detected by immunoblotting assay. **e,** Ribbon (left) and surface (right) structure of the C2 domain of Nedd4-1 with representation of XMU-MP-9-PAL-labelled residues (magenta) and residues critical for XMU-MP-9 binding (yellow). The structure of the C2 domain of Nedd4-1 is derived from PDB code 3B7Y. **f,** Ribbon structure of K-Ras^G12V^ (PDB: 8AZZ) with representation of XMU-MP-9-PAL-labelled residues (magenta) and residue critical for XMU-MP-9 binding (yellow). The magnified area shows a close-up view of these sites. **g,** F78A mutation of K-Ras^G12V^ prevents XMU-MP-9-promoted ubiquitination of K-Ras^G12V^. Nedd4-1, F/K-Ras^G12V^, and F/K-Ras^G12V/F78A^ purified from bacteria were subjected to an *in vitro* ubiquitination assay in the absence or presence of indicated concentrations of XMU-MP-9. **h,** F78A mutation decreases binding affinity of K-Ras^G12V^ to XMU-MP-9. Binding affinities of XMU-MP-9 to K-Ras^G12V^ and K-Ras^G12V/F78A^ were determined by MST assays and compared as fraction bound. Data are presented as the mean ±SD from three repeated experiments. **i,** K127A/F129A mutation of Nedd4-1 blocks XMU-MP-9-promoted ubiquitination of K-Ras^G12V^. F/K-Ras^G12V^, Nedd4-1, and Nedd4-1^K127A/F129A^ purified from bacteria were subjected to an *in vitro* ubiquitination assay in the absence or presence of indicated concentrations of XMU-MP-9. **j,** K127A/F129A mutation decreases binding affinity of Nedd4-1 to XMU-MP-9. Same as in (**h**) except Nedd4-1 and Nedd4-1^K127A/F129A^ were used.

We next exerted photo-affinity labeling technology^32^ to explore the binding pocket of XMU-MP-9 on K-Ras^G12V^ and Nedd4-1. We designed and synthesized XMU-MP-9-PAL (<u>p</u>hoto-<u>a</u>ffinity labeling), a diazirine derivative of XMU-MP-9, as a photo-affinity probe (Extended Data Fig. 3a). We performed mass spectrometry analysis to determine photo-affinity labeling sites in K-Ras^G12V^ and Nedd4-1 after they were individually incubated with XMU-MP-9-PAL and irradiated with UV light. By examining peptides with a mass shift +413.0810 Da, we found that V108, P109, and L110 of Nedd4-1 and I93, V112, and L113 of K-Ras^G12V^ could be labelled by XMU-MP-9-PAL (Extended Data Fig. 3b, c). The V108, P109, and L110 are located at a potential binding cleft in the C2 domain of Nedd4-1 (Fig. 3e), whereas the I93, V112, and L113 are located inside K-Ras^G12V^ (Fig. 3f). Interestingly, recent study suggested 8 novel allosteric sites in K-Ras^33^, and the I93, V112, and L113 are very close to one of these allosteric sites.

We analyzed the structure of K-Ras^G12V^ to look for potential amino acid residue(s) around this allosteric site critical for the binding of XMU-MP-9 and examined their effectiveness by site-directed mutagenesis. Among the mutations we have tested, F78A mutation significantly attenuated the effect of XMU-MP-9 on promoting Nedd4-1-mediated K-Ras^G12V^ ubiquitination and degradation (Fig. 3g and Extended Data Fig. 3d), and dramatically decreased the binding affinity of XMU-MP-9 to K-Ras^G12V^ (Fig. 3h). The enhancement of interaction between Nedd4-1 and K-Ras^G12V^ promoted by XMU-MP-9 was also abolished by F78A mutation of K-Ras^G12V^ (Extended Data Fig. 3e). We confirmed that F78A mutation did not affect K-Ras^G12V^ activity in triggering downstream MAP kinase pathway activation (Extended Data Fig. 3f), indicating that F78A mutation unlikely causes misfolding of K-Ras^G12V^. Using the same strategy, we identified that K127 and F129 of Nedd4-1 were critical for its activity to target K-Ras^G12V^ for ubiquitination and degradation upon XMU-MP-9 treatment (Fig. 3i and Extended Data Fig. 3g). Accordingly, K127A/F129A mutation significantly decreased the binding affinity of Nedd4-1 with XMU-MP-9 (Fig. 3j) and attenuated XMU-MP-9-induced enhancement of interaction between Nedd4-1 and K-Ras^G12V^ (Extended Data Fig. 3h). Meanwhile, K127A/F129A mutation did not affect Nedd4-1 to target wild-type K-Ras for degradation (Extended Data Fig. 3i), suggesting that K127 and F129 are not critical for Nedd4-1 folding and activity.

### XMU-MP-9 promotes ubiquitination and degradation of various K-Ras mutants

Besides G12V mutation, various mutations at residues 12, 13, and 61 of Ras such as G12D, G12C, G12S, G13V, G13D, and Q61H also occur frequently in various human cancers^7^. In addition, because Nedd4-1 is a general E3 ubiquitin ligase for controlling all three major forms of Ras (K-Ras, H-Ras, and N-Ras) protein levels^30^, we further compared the effects of XMU-MP-9 on Nedd4-1-mediated degradation of different mutants of the three forms of Ras. In agreement with our previous report^30^, overexpression of Nedd4-1 itself did not significantly affect the steady-state protein levels of these oncogenic mutants of K-Ras; however, treatment of XMU-MP-9 significantly induced decreases of the protein levels of all these K-Ras mutants. In contrast to its remarkable effects on inducing degradation of K-Ras mutants, XMU-MP-9 showed much less impact on degradation of H-Ras or N-Ras mutants (Extended Data Fig. 4a), indicating that XMU-MP-9 is a selective degrader for K-Ras mutants.

To compare the efficacies of XMU-MP-9 on promoting degradations of different K-Ras oncogenic mutants, we took advantage of endogenous Nedd4-1 in HEK293T cells to avoid potential variation of the protein levels of Nedd4-1 caused by overexpression. Meanwhile, we also generated Nedd4-1 knockout HEK293T cells for controls. XMU-MP-9 showed similar 50% effective concentrations (EC_50_ = 1.9 to 3.6 μM) to these K-Ras oncogenic mutants in wild-type HEK293T cells, whereas knockout of Nedd4-1 blocked the degradations of these K-Ras mutants induced by XMU-MP-9 (Fig. 4a and Extended Data Fig. 4b). Moreover, we confirmed that the XMU-MP-9-induced degradations of these K-Ras mutants were all via lysosomal pathway (Fig. 4b).

**Figure 4.**
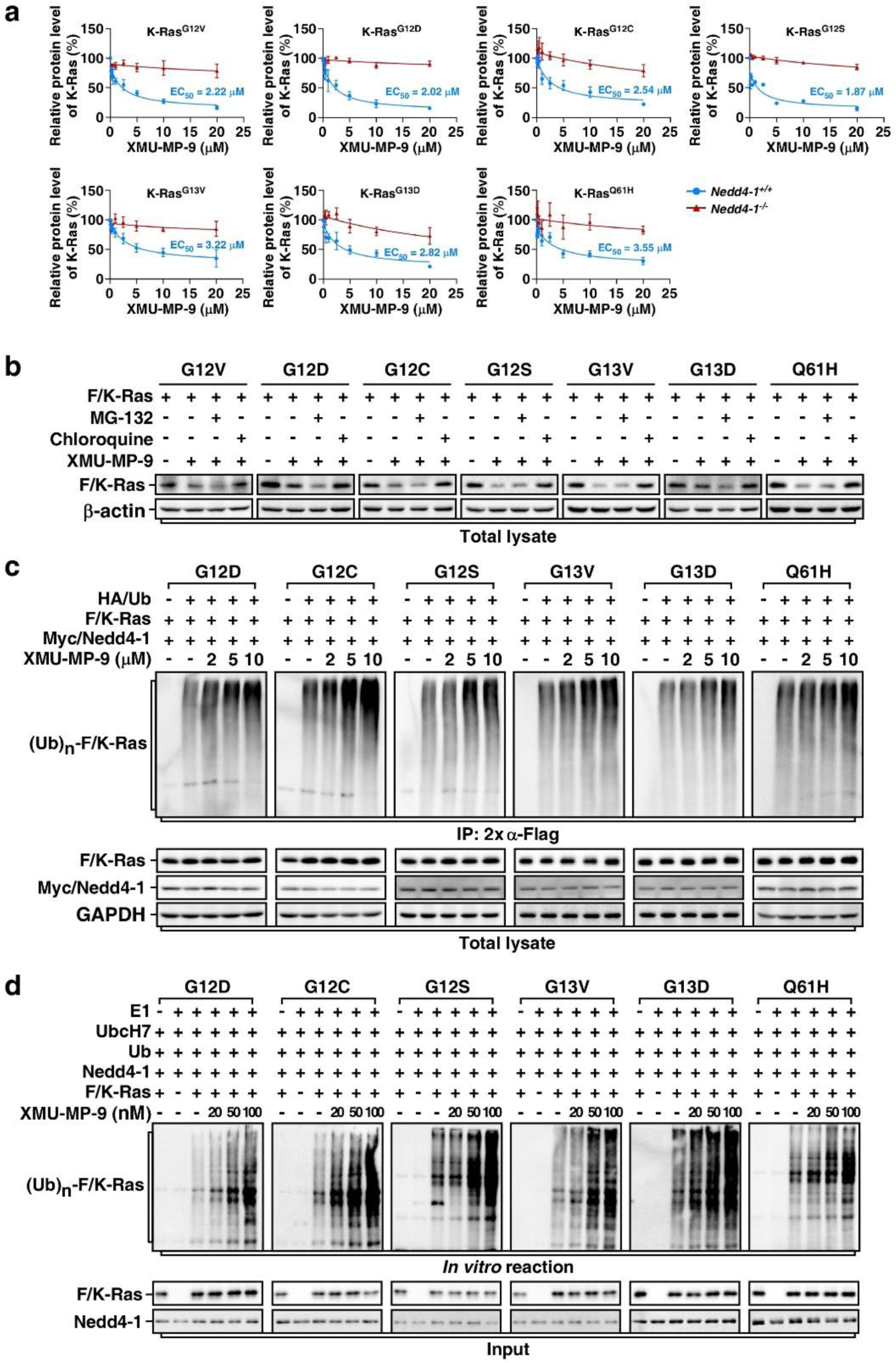
XMU-MP-9 has a general effect on promoting ubiquitination and degradation of various K-Ras oncogenic mutants. **a,** Quantification of relative protein levels of indicated K-Ras mutants treated 48 h with different concentrations of XMU-MP-9. Data were represented as mean ± SEM of 3 independent experiments. **b,** Degradation of K-Ras mutants induced by XMU-MP-9 is via lysosomal pathway. HEK293T cells transfected with indicated K-Ras mutants were treated with combinations of XMU-MP-9 (10 μM, 48 h) and proteasome inhibitor MG-132 (20 μM, 3 h) or lysosome inhibitor chloroquine (120 μM, 12 h) as indicated, and then subjected to immunoblotting assay. **c,** XMU-MP-9 promotes ubiquitination of various K-Ras mutants in cells. HEK293T cells transfected with indicated combinations of HA/Ub, Myc/Nedd4-1, and Flag-tagged K-Ras (F/K-Ras) mutants as indicated were pre-treated with chloroquine (120 μM, 1 h) and then treated 1 h with indicated concentrations of XMU-MP-9 before subjected to 2× anti-Flag IP followed by immunoblotting with anti-HA antibody to examine ubiquitin-conjugated F/K-Ras. Expression of Myc/Nedd4-1 and F/K-Ras were confirmed by immunoblotting the total cell lysates. **d,** XMU-MP-9 promotes ubiquitination of various K-Ras mutants *in vitro*. Bacterially expressed and purified Nedd4-1 and indicated F/K-Ras mutants were subjected to an *in vitro* ubiquitination assay in the presence or absence of indicated concentrations of XMU-MP-9.

We further verified the effects of XMU-MP-9 on promoting Nedd4-1-mediated ubiquitination of different K-Ras oncogenic mutants by executing ubiquitination assays in cells and *in vitro*. Similar as it did to K-Ras^G12V^, XMU-MP-9 dramatically promoted Nedd4-1-mediated ubiquitination of all the other K-Ras oncogenic mutants exogenously expressed in HEK293T cells (Fig. 4c). Additionally, *in vitro* ubiquitination assays using bacterially expressed Flag-tagged K-Ras mutants also confirmed that XMU-MP-9 could directly assist Nedd4-1 to ubiquitinate all these K-Ras mutants (Fig. 4d), demonstrating a general effect of XMU-MP-9 on promoting Nedd4-1-mediated ubiquitination of different K-Ras mutants. Hence, our study identified a general small-molecule degrader for different K-Ras oncogenic mutants by leveraging Nedd4-1, the natural E3 ubiquitin ligase for Ras proteins.

### XMU-MP-9 inhibits proliferation and anchorage-independent growth of K-Ras mutant harboring cancer cells

We next examined the effects of XMU-MP-9 on protein levels of endogenous K-Ras mutants and the downstream signaling. For this end, we treated various cancer cell lines harboring different oncogenic K-Ras mutations, including SW620 (colorectal, K-Ras^G12V^), AsPC-1 (pancreatic, K-Ras^G12D^), MIA PaCa-2 (pancreatic, K-Ras^G12C^), A549 (lung, K-Ras^G12S^), HCT 116 (colorectal, K-Ras^G13D^), and NCI-H460 (lung, K-Ras^Q61H^), with different doses of XMU-MP-9 and determined the endogenous K-Ras protein levels. In agreement with that XMU-MP-9 could promote Nedd4-1 to target various K-Ras mutants for ubiquitination (Fig. 4c, d), treatment of XMU-MP-9 dose-dependently induced decrease of endogenous K-Ras mutant protein levels in these cells. Accordingly, along with the decrease of K-Ras protein levels induced by treatment of XMU-MP-9, phosphorylation levels of B-Raf and MEK were markedly decreased (Fig. 5a).

**Figure 5.**
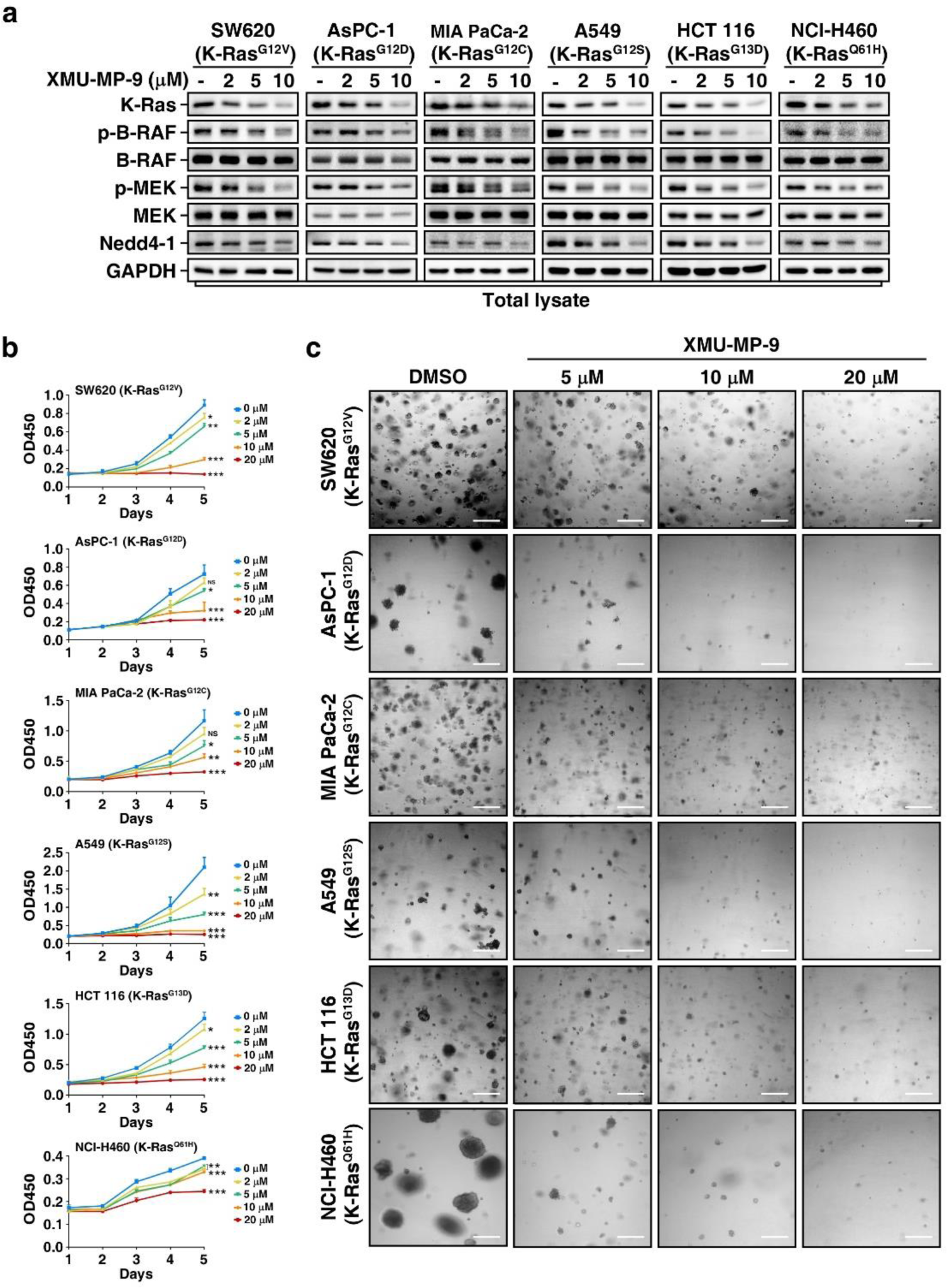
XMU-MP-9 induces degradation of endogenous K-Ras mutants and inhibits growth of K-Ras mutant harboring cancer cells. **a,** XMU-MP-9 promotes degradation of endogenous K-Ras mutants and inhibits its downstream signaling in K-Ras mutant harboring cancer cells. Different cancer cells harboring indicated K-Ras mutants were treated 48 h with indicated concentrations of XMU-MP-9 and then subjected to immunoblotting assay. **b,** XMU-MP-9 inhibits proliferations of K-Ras mutant harboring cancer cells in 2-D culture. Cancer cells harboring indicated K-Ras mutants were cultured with indicated concentrations of XMU-MP-9. Cell viabilities were determined and plotted as mean ± SD (n = 6). *P* values were determined by one-way ANOVA followed by Tukey test. * *p* < 0.05; ** *p* < 0.01; *** *p* < 0.001; NS, not significant. **c**, XMU-MP-9 suppresses anchorage-independent growth of K-Ras mutant harboring cancer cells. Cancer cells harboring indicated K-Ras mutants were seeded in 3-D soft agar and treated with indicated concentrations of XMU-MP-9 or DMSO as control for 2 weeks. Scale bar indicates 100 μm.

To probe the potential of XMU-MP-9 in K-Ras-driven cancer therapy, we first examined the growth inhibitory effects of XMU-MP-9 on these human cancer cells. As expected, proliferation of these cells in 2-D tissue culture were all substantially inhibited by treatment of XMU-MP-9 in a dose dependent manner (Fig. 5b). Meanwhile, growth of these cancer cells in 3-D soft agar culture were also dramatically inhibited by XMU-MP-9 (Fig. 5c), in line with the key role of Ras/Raf/MAPK signaling pathway in cancer cell proliferation and anchorage-independent growth^1^.

To confirm that XMU-MP-9 treatment-induced growth inhibition was through targeting Ras proteins, we treated HT-29, a human colorectal cancer cell line harboring constitutively active B-Raf^V600E^ mutant, with XMU-MP-9. As predicted, although the K-Ras protein levels were decreased by treatment of XMU-MP-9, phosphorylation levels of B-Raf^V600E^ and MEK were not changed (Extended Data Fig. 5a). In contrast to the remarkable inhibitory effects on both 2-D and 3-D growth of K-Ras mutant harboring cancer cells, XMU-MP-9 treatment showed a much less effect on inhibiting growth of HT-29 cells, especially in the 3-D soft agar assay (Extended Data Fig. 5b, c), suggesting that Ras/Raf/MAPK signaling pathway has a more pivotal role in promoting anchorage-independent growth than in promoting cell proliferation in 2-D culture. Thus, it is plausible that XMU-MP-9 indeed induces degradation of oncogenic K-Ras proteins to inhibit proliferation and anchorage-independent growth of cancer cells harboring oncogenic K-Ras mutations.

### XMU-MP-9 suppresses tumor development of K-Ras mutant harboring cancer cells *in vivo*

We next assessed the efficacy of XMU-MP-9 in suppressing tumor growth of K-Ras harboring cancer cells *in vivo*. To this end, we subcutaneously injected SW620 cells into nude mice to form xenograft tumors. XMU-MP-9 was then administered at a dose of 40 mg/kg once or twice daily via tail vein injection. Remarkably, treatment of XMU-MP-9 dose-dependently suppressed growth of xenograft tumors formed by SW620 cells (Fig. 6a-c). Accordingly, the protein levels of K-Ras^G12V^ were decreased in tumors treated with XMU-MP-9, and the phosphorylation levels of B-Raf and MEK were consequently attenuated, as examined by immunoblotting and immunohistochemistry assays (Fig. 6d, e). In line with the effects of XMU-MP-9 on growth of HT-29 cells (Extended Data Fig. 5b, c), treatment of XMU-MP-9 did not suppress growth of xenograft tumors formed by subcutaneously injected HT-29 cells into nude mice (Extended Data Fig. 6a-c). Accordingly, although the protein levels of K-Ras were decreased in XMU-MP-9 treated tumors, phosphorylation levels of B-Raf^V600E^ and MEK were not significantly changed (Extended Data Fig. 6d, e).

**Figure 6.**
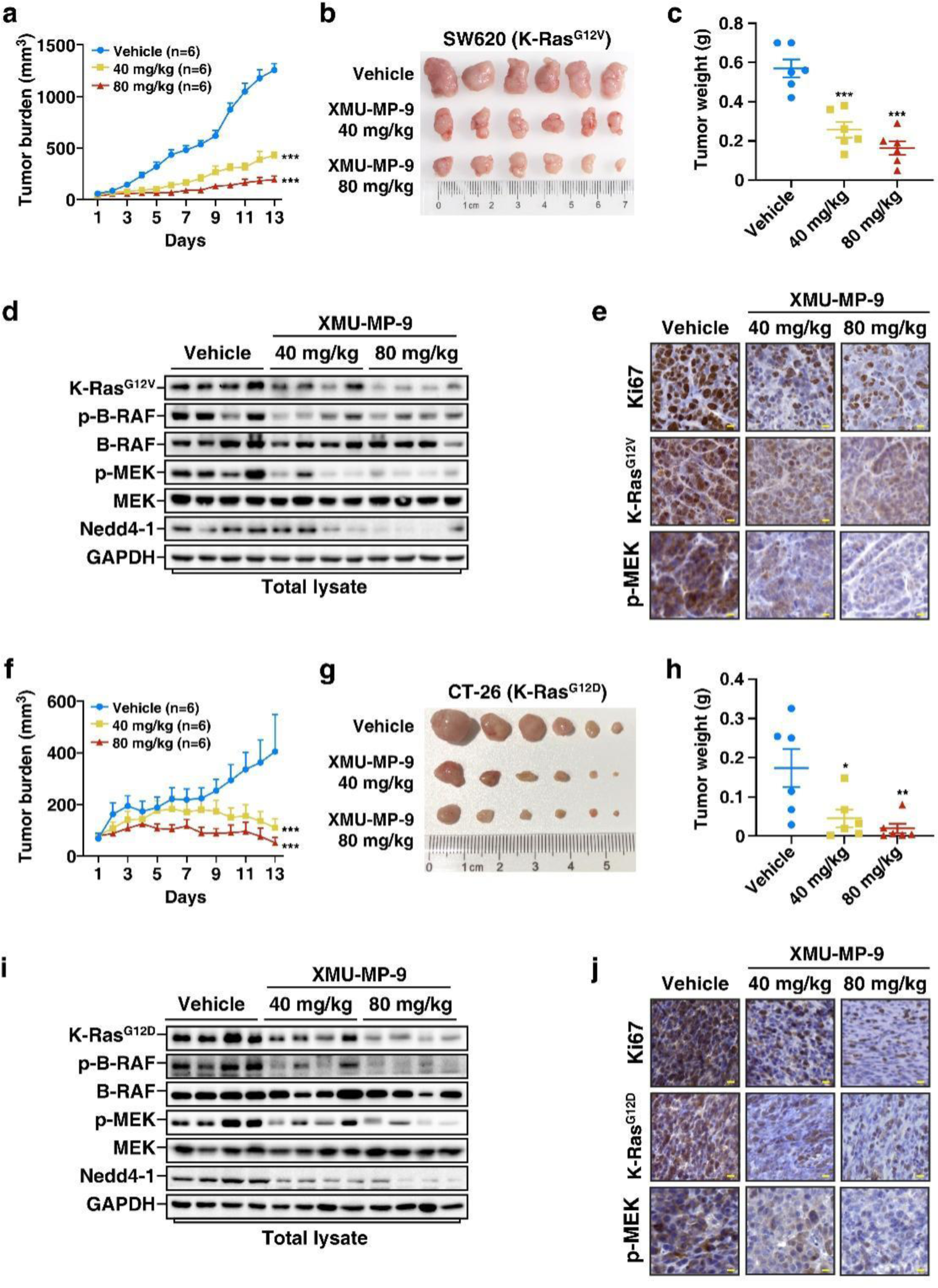
XMU-MP-9 suppresses growth of K-Ras mutant harboring cancer cells formed tumor in mice. **a-c,** XMU-MP-9 inhibits xenograft tumor growth of SW620 cells in nude mice. SW620 cells were subcutaneously injected into nude mice to form xenograft tumors. The tumor-bearing mice were given a daily dose of 40 mg/kg or 80 mg/kg of XMU-MP-9 (injected with a dose of 40 mg/kg once or twice a day via tail vein) or vehicle as control. Tumor volumes were measured and plotted as mean ±SEM (n = 6 animals per group). *P* values were determined by two-way ANOVA followed by Tukey test. *** *p* < 0.001 (**a**). The tumors were obtained 13 days after drug treatment by sacrificing the mice (**b**), weighted and plotted as mean ± SEM (n = 6 animals per group). *P* values were determined by one-way ANOVA followed by Tukey test. *** *p* < 0.001 (**c**). **d, e,** Treatment of XMU-MP-9 decreases K-Ras^G12V^ levels and downstream signaling in SW620 cells formed tumors. Representative tumors were subjected to immunoblotting (**d**) or IHC assay (**e**). Scale bar indicates 10 μm. **f-h,** XMU-MP-9 inhibits transplant tumor growth of CT-26 cells in BALB/c mice. CT-26 cells were subcutaneously injected into BALB/c mice to form transplant tumors. The tumor-bearing mice were treated and analyzed same as in (**a-c**). **i, j,** Treatment of XMU-MP-9 decreases K-Ras^G12D^ levels and downstream signaling in CT-26 cells formed tumors. Representative tumors were subjected to immunoblotting (**i**) or IHC assay (**j**). Scale bar indicates 10 μm.

Emerging evidences indicate that oncogenic mutations of K-Ras may lead to immune evasion in tumor microenvironment^34^. Therefore, we further evaluated the anti-tumor efficacy of XMU-MP-9 in immunocompetent mice. We subcutaneously injected CT-26, a mouse colorectal cancer cell line harboring K-Ras^G12D^, into BALB/c mice to generate transplanted tumors. Treatment of XMU-MP-9 even presented a more robust inhibitory effect on tumors formed by CT-26 cells in BALB/c mice than tumors formed by SW620 cells in nude mice (Fig. 6f-h). Similarly, the protein levels of K-Ras^G12D^ and the phosphorylation levels of B-Raf and MEK in CT-26 tumors treated with XMU-MP-9 were dramatically decreased (Fig. 6i, j). In addition, treatment of XMU-MP-9 did not show significant weight loss and obvious toxicity to heart, liver, spleen, lung, or kidney tissues of mice (Extended Data Fig. 6f, g). Hence, XMU-MP-9 demonstrated a good potential in treatment of K-Ras-driven cancers, which may lead to a new strategy for development of anticancer drugs.

## Discussion

Given the fact that Ras mutations occur so frequently in human cancers, directly targeting Ras mutants has always been an appealing approach for clinical therapy of Ras-driven cancers. While there have been significant breakthroughs in the development of allele-specific covalent inhibitors for K-Ras^G12C^ and the advancements in non-covalent inhibitor research for K-Ras^G12D^, designing small-molecule inhibitors for K-Ras mutants like K-Ras^G12V^, which lack specific active or charged side chains, remains a big challenge because there is no suitable binding pocket on the surface of these mutants^21,35^. Due to the same reason, PROTACs for targeted degradation of K-Ras mutants has only been designed for K-Ras^G12C^ based on the existing inhibitors^36,37^. Thus, it is highly desirable to develop innovative strategies for targeting these Ras mutants commonly found in human cancers.

Our previous study demonstrated that Nedd4-1 is a natural E3 ubiquitin ligase for Ras proteins. Although oncogenic Ras mutants are resistant to Nedd4-1-mediated ubiquitination, they still can bind Nedd4-1^30^. Based on this finding, we utilized Nedd4-1, instead of hijacking an irrelevant E3 ubiquitin ligase that is usually used in PROTAC or molecular glue degrader design, to seek degraders for K-Ras mutants. This strategy allowed compounds to work on the pre-existing Nedd4-1 and K-Ras interaction and led to successful identification of a general small-molecule degrader for all major K-Ras mutants. Moreover, Nedd4-1 is also an E3 ubiquitin ligase for H-Ras and N-Ras^30^. It is possible to obtain small-molecule degraders for H-Ras or N-Ras mutants as well using the same strategy. Hence, our study presents a promising approach to achieve small-molecule degraders for all kinds of Ras mutants.

Because there is no specific active or charged side group available on Ras mutants like Ras^G12V^, it will be extremely hard to specifically target these Ras mutants without affecting wild-type Ras in normal tissues. This raises safety concerns for using inhibitors to target these Ras mutants in patients. Our previous study indicated that Ras signaling is required for transcription of Nedd4-1. The negative feedback regulatory loop between Nedd4-1-mediated Ras degradation and Ras signaling-regulated Nedd4-1 transcription plays a key role in maintaining homeostasis of Ras signaling in normal cells. Oncogenic mutations of Ras block Nedd4-1-mediated Ras degradation, resulting in uncontrolled Ras signaling and, as a consequence, upregulated Nedd4-1 levels^30^. In fact, Nedd4-1 was found frequently upregulated in various types of human cancers and showed a positive correlation with Ras levels^30,38^, thus providing a therapeutic window for treatment of Ras-driven cancers with Nedd4-1-dependent Ras degrader. Indeed, treatment of XMU-MP-9 promoted degradation of K-Ras mutants and consequently decreased endogenous levels of Nedd4-1 (Fig. 1h, Fig. 5a and Fig. 6d, i), suggesting that Nedd4-1-dependent Ras degradation will be restrained when Ras signaling is low, which may considerably lessen its potential toxic effects to normal tissues. As a support, our experiment revealed that treatment of XMU-MP-9 did not show obvious toxicity to the treated mice (Extended Data Fig. 6f, g).

In comparison with molecular glue degraders, PROTACs generally have poorer drug-like properties due to their relatively larger sizes^26,39^. Although functions as a bifunctional compound, XMU-MP-9 does not contain a flexible linker and has a relatively small molecular weight. Rather than induces a non-native interaction, XMU-MP-9 stabilizes the pre-existing interaction between Nedd4-1 and K-Ras mutants. More interestingly, it looks like that XMU-MP-9 not only enhances the interaction between Nedd4-1 and K-Ras^G12V^ (Fig. 3c, d), but also induces a conformational change of K-Ras^G12V^ to expose its K128 to Nedd4-1. In line with this, the binding site of XMU-MP-9 identified in K-Ras^G12V^ is close to an allosteric site that was recently reported by Weng et al.^33^, and K128 is also around this allosteric site. Binding of XMU-MP-9 might stabilize this allosteric conformation that allows Nedd4-1 to access to K128 of K-Ras^G12V^ (Fig. 2f, g). Therefore, in addition to its binding property, XMU-MP-9 may also act as a molecular wedge for stabilizing the transit conformation of K-Ras^G12V^. It will be of great interest to apply the conformation inducing/stabilizing function to further small-molecule degrader design for drug development.

## Materials and methods

### Antibodies and chemical reagents

Mouse anti-GAPDH (sc-32233), anti-β-actin (sc-47778), anti-Ub (sc-8017), and anti-Myc (SC-40) antibodies were purchased from Santa Cruz Biotechnology (Santa Cruz, CA, USA); rabbit anti-Ki67 (12202T), anti-Nedd4-1 (3607), anti-phospho-B-RAF (2696), anti-B-RAF (14814), anti-phospho-MEK1/2 (9154), and anti-MEK1/2 (9122) antibodies were purchased from Cell Signaling Technology; rabbit anti-K-Ras (BS6231) antibody was purchased from Bioworld; rabbit anti-K-Ras (A1190) antibody was purchased from ABclonal; mouse anti-FLAG M2 antibody (F1804) was from Sigma-Aldrich; and rat anti-HA antibody (11867431001) was from Roche. Cycloheximide and lysosome inhibitor chloroquine were from Sigma-Aldrich. Proteasome inhibitor MG-132 was from Boston Biochem. Cell Counting Kit-8 (CCK-8) was from MedChemExpress. FastKing RT Kit and SuperReal PreMix Plus (SYBR Green) were from TIANGEN.

### DNA constructs

The complementary DNA (cDNA) of human Nedd4-1 was a generous gift from Dr. W. Sundquist. Human cDNAs of K-Ras, H-Ras, and N-Ras were cloned from HEK293T cells using RT-PCR, and various mutations of Ras were introduced by PCR-based site-directed mutagenesis. Cloning for protein expression in mammalian cells was carried out using a modified pCMV5 vector for transfection, pBOBI vector for lentivirus infection. pGEX-4T-1 and pProEX were used for bacterial expression of proteins.

### Cell culture, transfection, and lentivirus infection

Human embryonic kidney HEK293T (CRL-3216), human colon cancer SW620 (CCL-227), HCT 116 (CCL-247), human pancreas cancer AsPC-1 (CRL-1682), MIA PaCa-2 (CRM-CRL-1420), human lung cancer NCI-H460 (HTB-177), A549 (CRM-CCL-185), and mouse colon cancer CT-26 (CRL-2638) cell lines were obtained from ATCC. Human colon cancer HT-29 (SCSP-5032) cell line was purchased from the Cell Bank of the Chinese Academy of Sciences (Shanghai). SW620 cells were cultured in Leibovitz’s L-15 medium (Basal Media); HEK293T, A549 cells were cultured in high-glucose Dulbecco’s modified Eagle medium (Gibco); MIA PaCa-2 was cultured in high-glucose Dulbecco’s modified Eagle medium (Gibco) added with 1 mM sodium pyruvate; AsPC-1, NCI-H460, and CT-26 were cultured in RPMI medium 1640 (Gibco); HCT 116 and HT-29 cells were cultured in McCoy’s 5A medium (Sigma-Aldrich), all supplemented with 10% (v/v) fetal bovine serum (Gibco) and 100 U/ml streptomycin and penicillin (Millipore). All cells were cultured at 37°C in a humidified incubator with 5% CO_2_ except SW620 with 0% CO_2_. The cell lines were routinely tested and confirmed to be negative for mycoplasma.

Polyethylenimine (PEI) was used for transient transfection of HEK293T cells as previously described^40^. Briefly, mixtures of PEI and plasmids (3:1, w/w) were added to cell cultures and incubated for 6 h before replaced with fresh medium. Recombinant lentivirus used to infect SW620 cells for protein expression was generated by the ViraPower Lentiviral Expression System (Invitrogen) according to the manufacturer’s instruction.

### Generation of Nedd4-1^-/-^ cell lines

For gene silencing of Nedd4-1 in SW620 and HEK293T cells, design of sgRNAs and off-target sites prediction were performed by using the Zhang laboratory CRISPR design tool (https://zlab.bio/guide-design-resources). The sequences for human Nedd4-1 gRNA-1 and gRNA-2 are 5’-TCAATCATGGGTAGGCAGAC-3’ and 5’-TCCAAGTTACTTGACGGTGG-3’, respectively. pLentiCRISPRv2 vector was used to express sgRNAs and the pool of Nedd4-1^-/-^ cells were selected by treatment with puromycin (2 μg/ml) for 7 days. The efficiency of Nedd4-1 knockout was assessed by quantitative reverse transcription-PCR (qRT-PCR) and immunoblotting assays.

### Cell viability and 3-D soft agar assays

Cell growth rate was determined by counting the cell numbers in triplicate daily after seeding into 96-well plate using a CCK-8 kit following the manufacturer’s instruction. Briefly, the culture plates were incubated with CCK-8 solution (10 µl/well) for 1 h, and then measured the absorbance at 450 nm using a microplate reader (Tecan Spark).

For 3-D soft agar assay, cells in 500 μl of 0.25% (w/v) Noble agar (BD) with complete medium were seeded to each well of 24-well plates precoated with equal volume of 1% (w/v) Noble agar. Additional 300 μl of overlay medium was added after cell plating and the overlay medium was changed every 3 days. The cells were cultured for 2 weeks and then subjected to photographing using a Nikon eclipse Ti-U inverted microscope to examine colony formation in soft agar.

### Microscale thermophoresis (MST) assay

Target proteins were fluorescently labeled using the Protein Labeling Kit RED-NHS (NanoTemper Technologies). Labeling reaction was performed following the manufacturer’s instructions with 10 μM protein. Labeled protein was adjusted to 20 nM with PBS buffer supplemented with 0.02 % Tween 20. The ligand (XMU-MP-9 or Nedd4-1) was dissolved in PBS buffer supplemented with 10% DMSO and serially diluted using the same buffer. Additional XMU-MP-9 was supplied as required for examining Nedd4-1 and K-Ras^G12V^ interaction. For the measurement, fluorescently labeled protein and ligand were mixed in 1:1 volume and then incubated for 5 minutes at room temperature before loaded into Monolith NT.115 Premium Capillaries (NanoTemper Technologies). MST was measured using a Monolith NT.115 instrument (NanoTemper Technologies) at 25°C with parameters adjusted to auto-detected LED power and medium MST power. Data of three independently pipetted measurements were analyzed using the MO Affinity Analysis software (NanoTemper Technologies).

### Transplant tumor formation assay

For generation of xenograft tumors of human colorectal SW620 and HT-29 cells in nude mice and transplant tumors of mouse colorectal CT-26 cells in BALB/c mice, 1-2×10^6^ cells suspended in 100 μl PBS were subcutaneously injected in nude mice or BALB/c mice. The growth of tumors was measured with digital calipers every day for 10-14 days after the tumors were visible. Nude mice and BALB/c mice were purchased from and housed in the Laboratory Animal Center of Xiamen University (China). All procedures involving mice for transplant tumor growth assays were approved by the Institutional Animal Care and Use Committee of Xiamen University.

### Immunoprecipitation, immunoblotting, and nickel (Ni) pull-down assays

Immunoprecipitation and immunoblotting were carried out as previously described^41^. Briefly, cells were lysed on ice with TNTE lysis buffer (50 mM tris-HCl pH 7.5, 150 mM NaCl, 1 mM EDTA, 0.5% Triton X-100, 10 μg/ml pepstatin A, 10 μg/ml leupeptin, and 1 mM phenylmethylsulfonyl fluoride) and cell lysates were collected by centrifugation at 20,000 g at 4°C for immunoprecipitation or immunoblotting assays with appropriate antibodies. For Ni pull-down assay, bacterially expressed GST/Nedd4-1 was purified using glutathione Sepharose beads in TNTE lysis buffer and free Nedd4-1 protein was obtained by tobacco etch virus (TEV) protease cleavage. His-tagged K-Ras^G12V^ was purified using Ni^2+^-NTA-agarose beads for Ni pull-down assay.

### Ubiquitination assay

For ubiquitination assay of Flag-tagged K-Ras in cells, cell lysates were subjected to anti-Flag immunoprecipitation, eluted by boiling in 1% SDS for 5 min, diluted 10 times in TNTE lysis buffer, and then re-immunoprecipitated with anti-Flag (2×IP). The ubiquitin-conjugated proteins were detected by immunoblotting HA-tagged ubiquitin (HA/Ub) with anti-HA antibody. For ubiquitination assay of endogenous K-Ras, cell lysates were subjected to anti-K-Ras immunoprecipitation flowed by immunoblotting. For *in vitro* ubiquitination assays, bacterially expressed and purified F/K-Ras and Nedd4-1 proteins were applied to *in vitro* ubiquitination reaction as previously described^41^. Reactions were stopped by boiling in 1% SDS for 5 min, and then diluted 10 times in TNTE lysis buffer before subjected to anti-Flag IP, followed by immunoblotting ubiquitin with anti-Ub antibody to examine ubiquitin-conjugated F/K-Ras proteins.

### Immunohistochemistry assay

For immunohistochemistry assay, tumors and tissues were fixed in 4% (v/v) paraformaldehyde (PFA), embedded in paraffin, sectioned at 3 μm, deparaffinized, and rehydrated. Sections were stained with hematoxylin and eosin (H&E) to examine the morphology. Immunohistochemical staining was performed using the UltraSensitiveTM SP kit (MXB, KIT-9720) with appropriate primary antibodies overnight at 4°C. Chromogenic revelation was performed with DAB kit (MXB, DAB-2031). Images were obtained using Zeiss AxioScan7.

### Photo-Affinity Labeling

Purified protein (K-Ras^G12V^ or Nedd4-1) was incubated with 10 μM XMU-MP-9-PAL for 30 min at 4°C in PBS buffer and then photo-crosslinked by 365 nm UV irradiation for 20 min on ice. After photo-affinity labeling, protein sample was mixed with an equal volume of 2× SDS sample loading buffer followed by SDS gel electrophoresis. Protein band was excised from the gel and subjected to LC-MS/MS analysis.

### Quantitative real-time PCR

Quantitative real-time PCR (qRT-PCR) was performed using SuperReal PreMix Plus (SYBR Green) (TIANGEN) according to the manufacturer’s protocol on a Mastercycler EP gradient S RealPlex2 (Eppendorf). Total RNAs for qRT-PCR were extracted from cells using TRIzol (Invitrogen), and cDNA was synthesized with FastKing RT kit (TIANGEN). The relative changes of gene expression were determined using the 2^-ΔΔCt^ method and normalized to GAPDH. The primers for human K-Ras are 5’-GAGTACAGTGCAATGAGGGAC-3’ (forward) and 5’-CCTGAGCCTGTTTTGTGTCTAC-3’ (reverse); for human Nedd4-1 are 5’-CAGGCCCTCAATCACAAGC-3’ (forward) and 5’-AGGCCCTAGATCATTGGAAGT-3’ (reverse); for human GAPDH are 5’-CATGAGAAGTATGACAACAGCCT-3’ (forward) and 5’-AGTCCTTCCACGATACCAAAGT-3’ (reverse).

### Statistics and reproducibility

GraphPad Prism 8 software was used to analyze all quantitative data. The data are represented as mean ± SEM or mean ± SD calculated using GraphPad. Significance was tested using unpaired two-tailed Student’s t test, one-way ANOVA with Tukey test and two-way ANOVA with Tukey test. Protein bands in immunoblotting assay were visualized by Sagecreation MiniChemi system and analyzed with Lane 1D software. *P* < 0.05 was considered a statistically significant difference (* *p* < 0.05, ** *p* < 0.01, and *** *p* < 0.001). All data are representative of at least three independent experiments unless otherwise specified.

## Data availability

All data supporting the findings of this study are available within the article and its Extended Data Information files or from the corresponding author upon request.

## Acknowledgements

This work was supported by the National Natural Science Foundation of China (82273037 to H.-R. Wang; 22025702, 82021003, 82151211 and 92253303 to X. Deng; 32070761 to T. Zeng), the National Key R&D Program (2022YFC2804100 to X. Deng), the Natural Science Foundation of Fujian Province (2022J05006 to T. Zeng), the Fundamental Research Funds for the Chinese Central Universities (20720230070 to T. Zeng), the Project “111” sponsored by the State Bureau of Foreign Experts and Ministry of Education of China (#BP2018017), and the New Cornerstone Science Foundation through the XPLORER PRIZE (To X. Deng).

## Author contribution

T. Zeng and T. Jiang conducted the experiments and analyzed the data. B. Zhang, T. Zhang, and X. Deng conceived and performed small-molecule degrader design and chemical synthesis. W. Dai and Y. Li helped with the animal experiments. X. Yin performed molecular biology experiments and protein purification. C. Wu performed MST assay. Y. Wu performed mass spectrometry. X. Chi helped for binding site identification. H.-R. Wang conceived the idea, supervised the study, and wrote the manuscript.

---

**Extended Data Figure 1.**
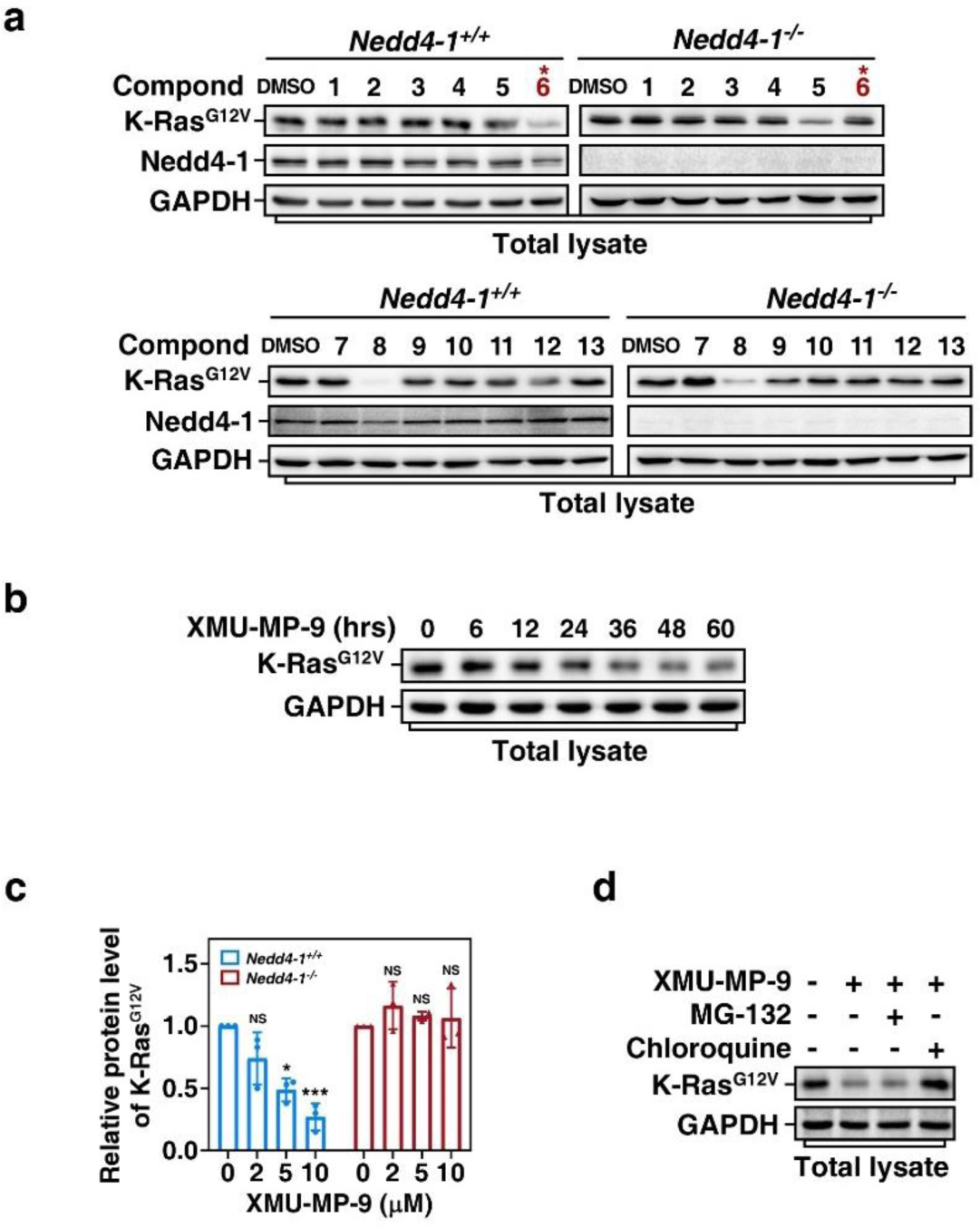
Screening for Nedd4-1-dependent K-Ras^G12V^ degrader. **a,** Screening for compounds able to promote Nedd4-1-dependent K-Ras^G12V^ degradation. Nedd4-1^+/+^ and Nedd4-1^-/-^ SW620 cells were treated 48 h with 10 μM compounds picked from the screen and then subjected to immunoblotting assay. The compound marked with red number (XMU-MP-9) was selected. **b**, XMU-MP-9 treatment down-regulates endogenous K-Ras^G12V^ protein levels. SW620 cells were treated with XMU-MP-9 (10 µM) for the indicated time and then subjected to immunoblotting assay. **c**, XMU-MP-9 dose-dependently induces decrease of endogenous K-Ras^G12V^ protein levels in Nedd4-1^+/+^ but not Nedd4-1^-/-^ SW620 cells. The protein levels of K-Ras^G12V^ were determined and plotted as mean ± SD of 3 independent experiments. *P* values were determined by one-way ANOVA followed by Tukey test. * *p* < 0.05; ** *p* < 0.01; *** *p* < 0.001; NS, not significant. **d**, XMU-MP-9 induces degradation of K-Ras^G12V^ via lysosomal pathway. SW620 cells treated as indicated combinations of XMU-MP-9 (10 μM, 48 h) and proteasome inhibitor MG-132 (20 μM, 3 h) or lysosome inhibitor chloroquine (120 μM, 12 h) were subjected to immunoblotting assay.

**Extended Data Figure 2.**
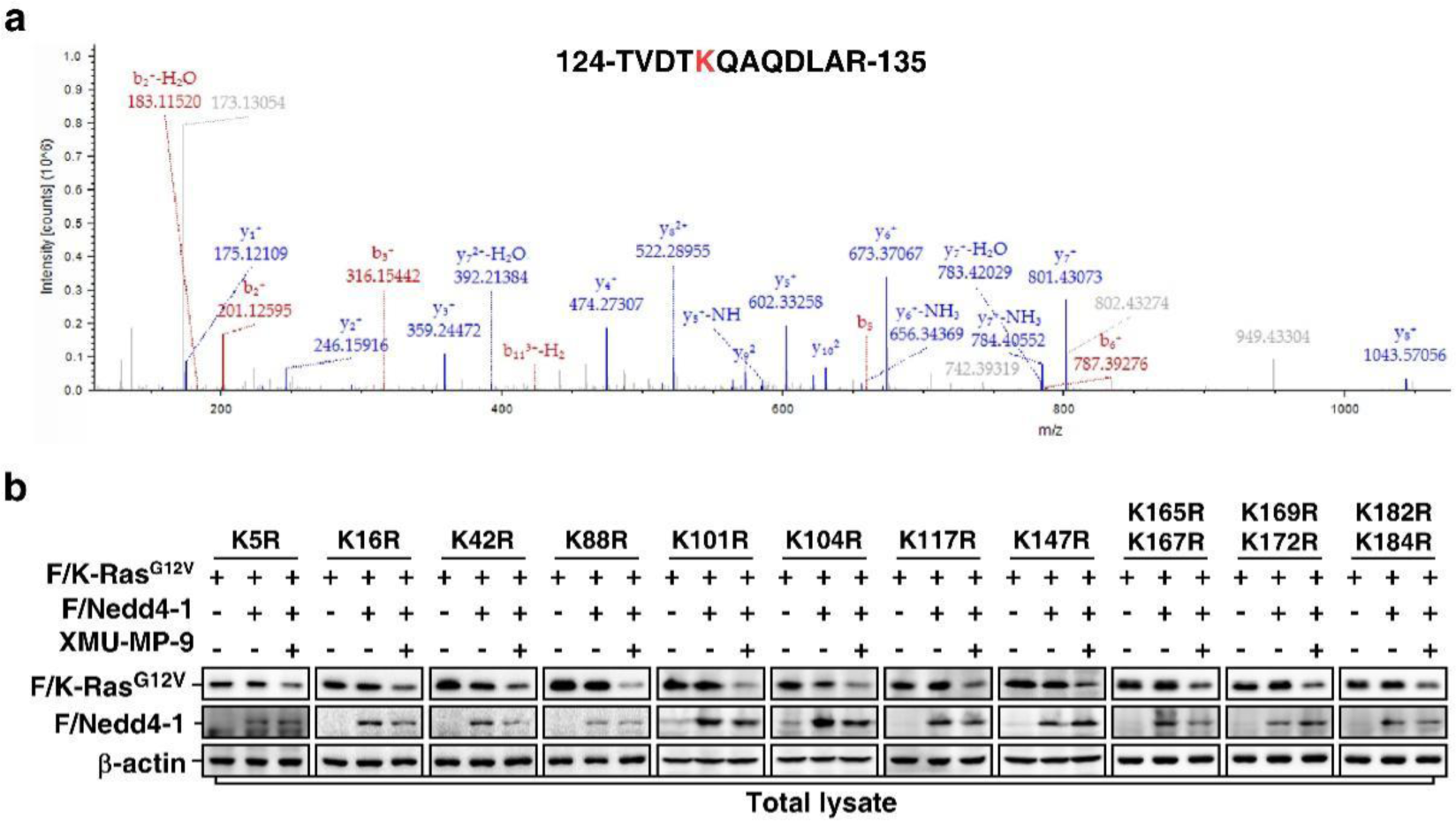
Determination of ubiquitination site of K-Ras^G12V^. **a**, Bacterially expressed K-Ras^G12V^ was applied to an *in vitro* ubiquitination reaction in the presence of 10 μM XMU-MP-9 and then subjected to mass spectrometry analysis to determine ubiquitination site(s) of K-Ras^G12V^. **b**, Effects of different Lys-to-Arg mutations on XMU-MP-9-induced K-Ras^G12V^ degradation. HEK293T cells transfected with F/Nedd4-1 and indicated Lys-to-Arg mutants of F/K-Ras^G12V^ were treated with or without XMU-MP-9 (10 μM, 48 h) and then subjected to immunoblotting assay.

**Extended Data Figure 3.**
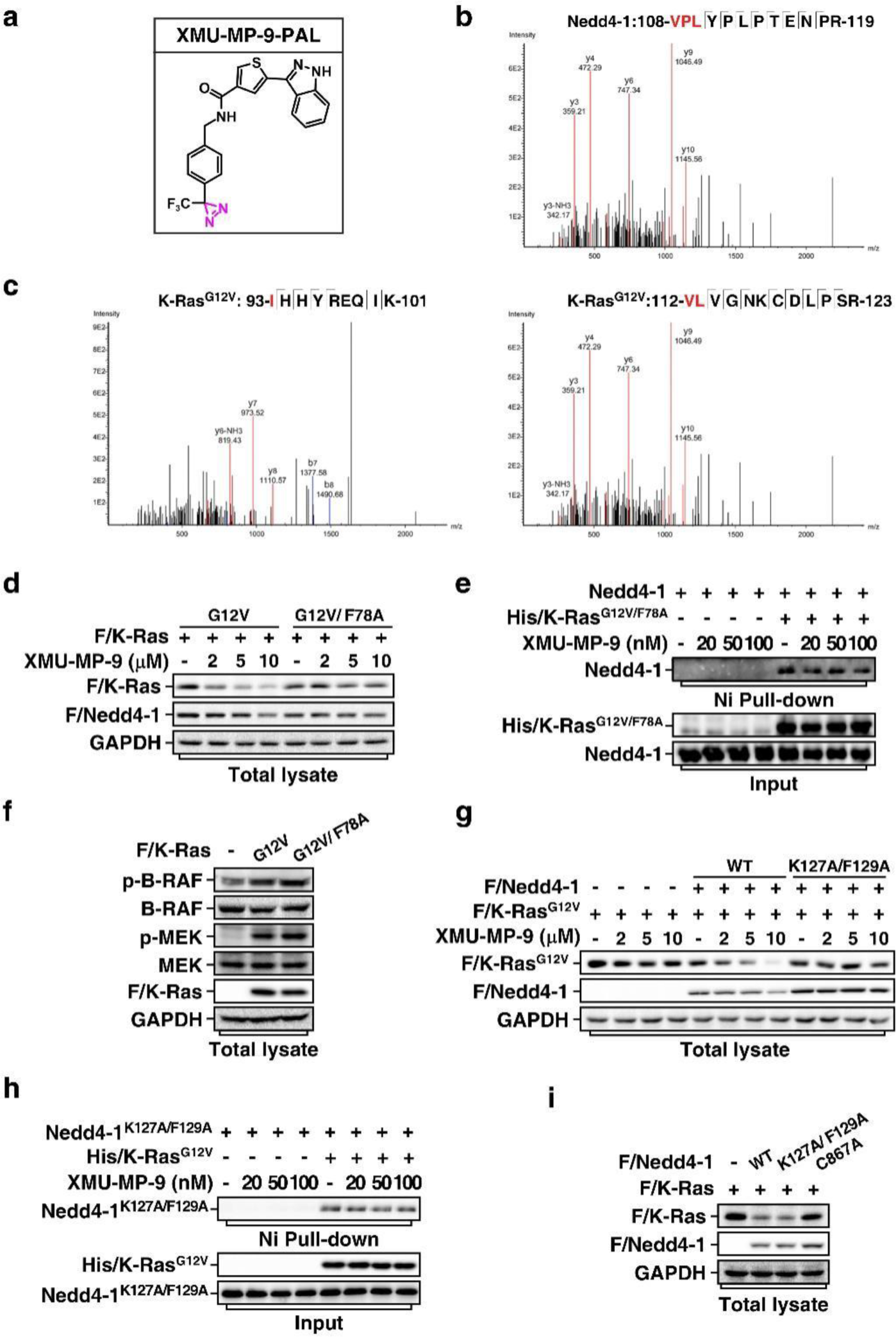
Identification of XMU-MP-9 binding sites on K-Ras^G12V^ and Nedd4-1. **a,** Chemical structure of XMU-MP-9-PAL, the photo-affinity probe derivate of XMU-MP-9. **b,** Identification of XMU-MP-9-PAL-labelled amino acid residues in Nedd4-1. Bacterially expressed and purified Nedd4-1 was photo-labelled by XMU-MP-9-PAL and subjected to mass spectrometry to determine the labelled amino acid residues. **c,** Identification of XMU-MP-9-PAL-labelled residues in K-Ras^G12V^. Same as in (**b**) except K-Ras^G12V^ was used. **d,** F78A mutation of K-Ras^G12V^ attenuates XMU-MP-9-induced degradation of K-Ras^G12V^ via Nedd4-1. HEK293T cells transfected with F/Nedd4-1 and F/K-Ras^G12V^ or F/K-Ras^G12V/F78A^ as indicated were treated with 48 h with indicated concentrations of XMU-MP-9 and then subjected to immunoblotting assay. **e,** F78A mutation of K-Ras^G12V^ prevents XMU-MP-9 to enhance the interaction between K-Ras^G12V^ and Nedd4-1. Bacterially expressed and purified His-tagged K-Ras^G12V, F78A^ (His/K-Ras^G12V/F78A^) and Nedd4-1 were subjected to Ni affinity pull-down assay. **f,** F78A mutation does not affect activity of K-Ras^G12V^. HEK293T cells transfected with K-Ras^G12V^ or F/K-Ras^G12V/F78A^ were subjected to immunoblotting assay. **g,** K127A/F129A mutation of Nedd4-1 disrupts XMU-MP-9-induced degradation of K-Ras^G12V^. Nedd4-1^-/-^ HEK293T cells transfected with F/K-Ras^G12V^ and F/Nedd4-1 or F/Nedd4-1^K127A/F129A^ as indicated were treated 48 h with indicated concentrations of XMU-MP-9 and then subjected to immunoblotting assay. **h,** K127A/F129A mutation of Nedd4-1 blocks XMU-MP-9-enhanced interaction between K-Ras^G12V^ and Nedd4-1. Same as in (**e**) except His/K-Ras^G12V^ and Nedd4-1^K127A/F129A^ were used. **i,** K127A/F129A mutation does not affect activity of Nedd4-1 in targeting wild-type K-Ras for degradation. HEK293T cells transfected with indicated combinations of wild-type F/K-Ras, wild-type F/Nedd4-1, F/Nedd4-1^K127A/F129A^, and catalytic inactive F/Nedd4-1^C867A^ were subjected to immunoblotting assay.

**Extended Data Figure 4.**
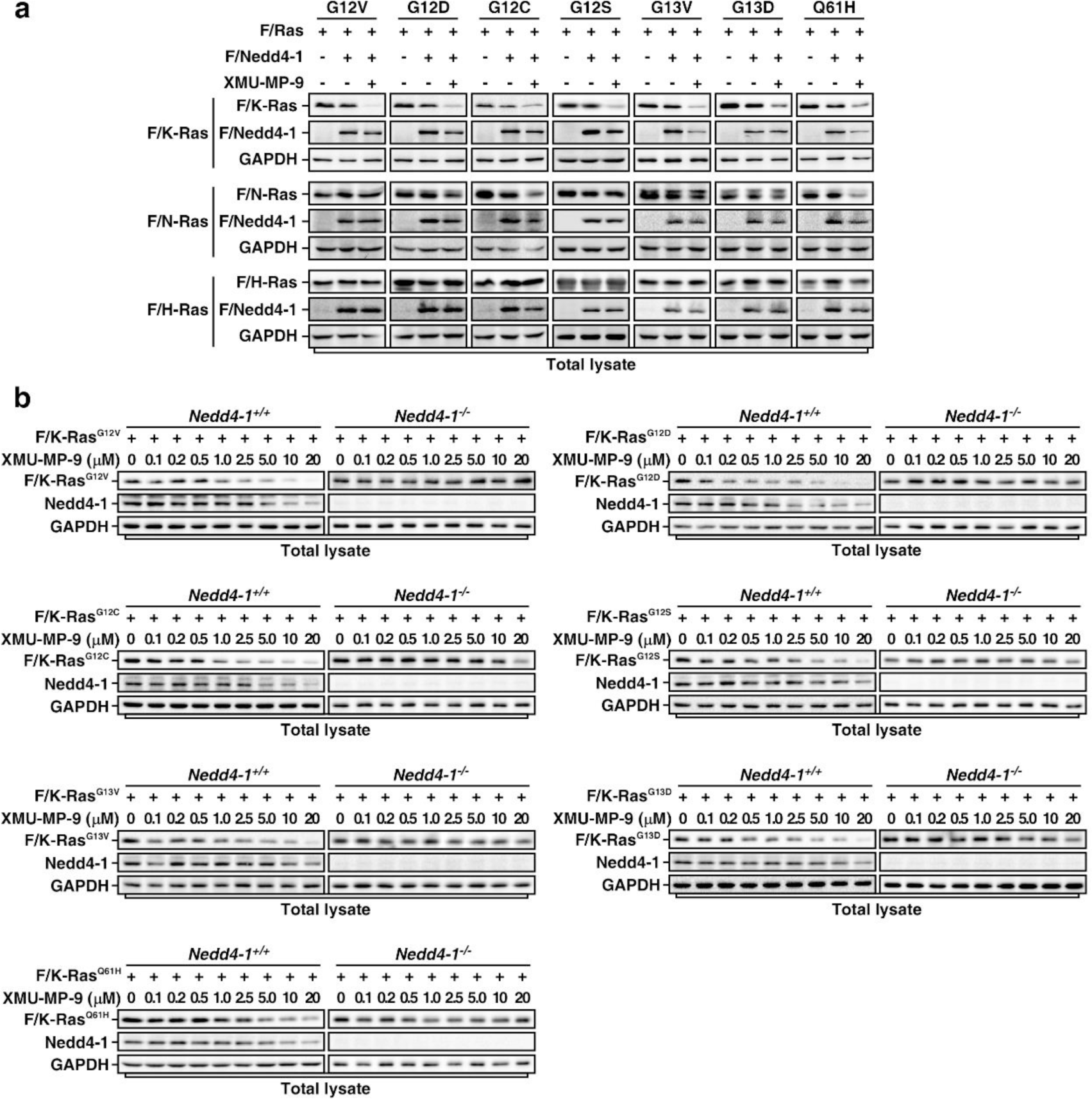
XMU-MP-9 promotes degradation of various K-Ras oncogenic mutants. **a**, XMU-MP-9 has a more potent effect on inducing degradation of K-Ras mutants than N-Ras or H-Ras mutants. HEK293T cells transfected with F/Nedd4-1 and indicated mutants of F/K-Ras, Flag-tagged N-Ras (F/N-Ras), or Flag-tagged H-Ras (F/H-Ras) were treated 48 h with or without 10 μM XMU-MP-9 and then subjected to immunoblotting assay. **b,** XMU-MP-9 promotes degradation of K-Ras oncogenic mutants in a dose-dependent manner in the presence of Nedd4-1. Nedd4-1^+/+^ or Nedd4-1^-/-^ HEK293T cells transfected with indicated Flag-tagged K-Ras mutants were treated 48 h with indicated concentrations of XMU-MP-9 and then subjected to immunoblotting assay.

**Extended Data Figure 5.**
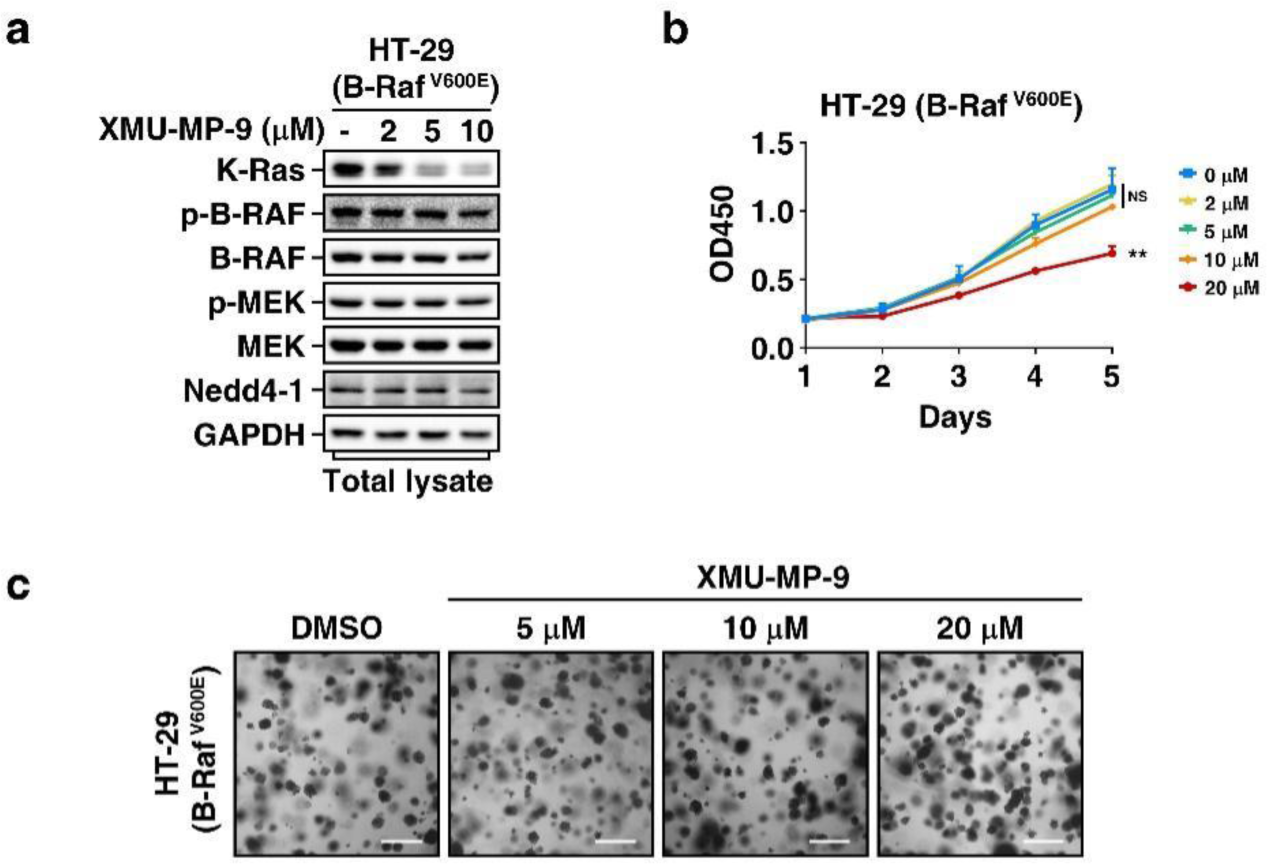
Effects of XMU-MP-9 on growth of B-Raf mutant harboring cells. **a**, XMU-MP-9 promotes K-Ras degradation but does not affect downstream Raf/MEK signaling in B-Raf^V600E^ harboring cells. B-Raf^V600E^ harboring HT-29 cells were treated 48 h with indicated concentrations of XMU-MP-9 and then subjected to immunoblotting assay. **b,** XMU-MP-9 has a minor effect on inhibiting proliferation of HT-29 in 2-D culture. Viabilities of HT-29 cells cultured with indicated concentrations of XMU-MP-9 were determined and plotted as mean ± SD (n = 6). *P* values were determined by one-way ANOVA followed by Tukey test. ** *p* < 0.01; NS, not significant. **c,** XMU-MP-9 does not suppress anchorage-independent growth of HT-29 cells. HT-29 cells seeded in 3-D soft agar were treated with indicated concentrations of XMU-MP-9 or DMSO for 2 weeks. Scale bar indicates 100 μm.

**Extended Data Figure 6.**
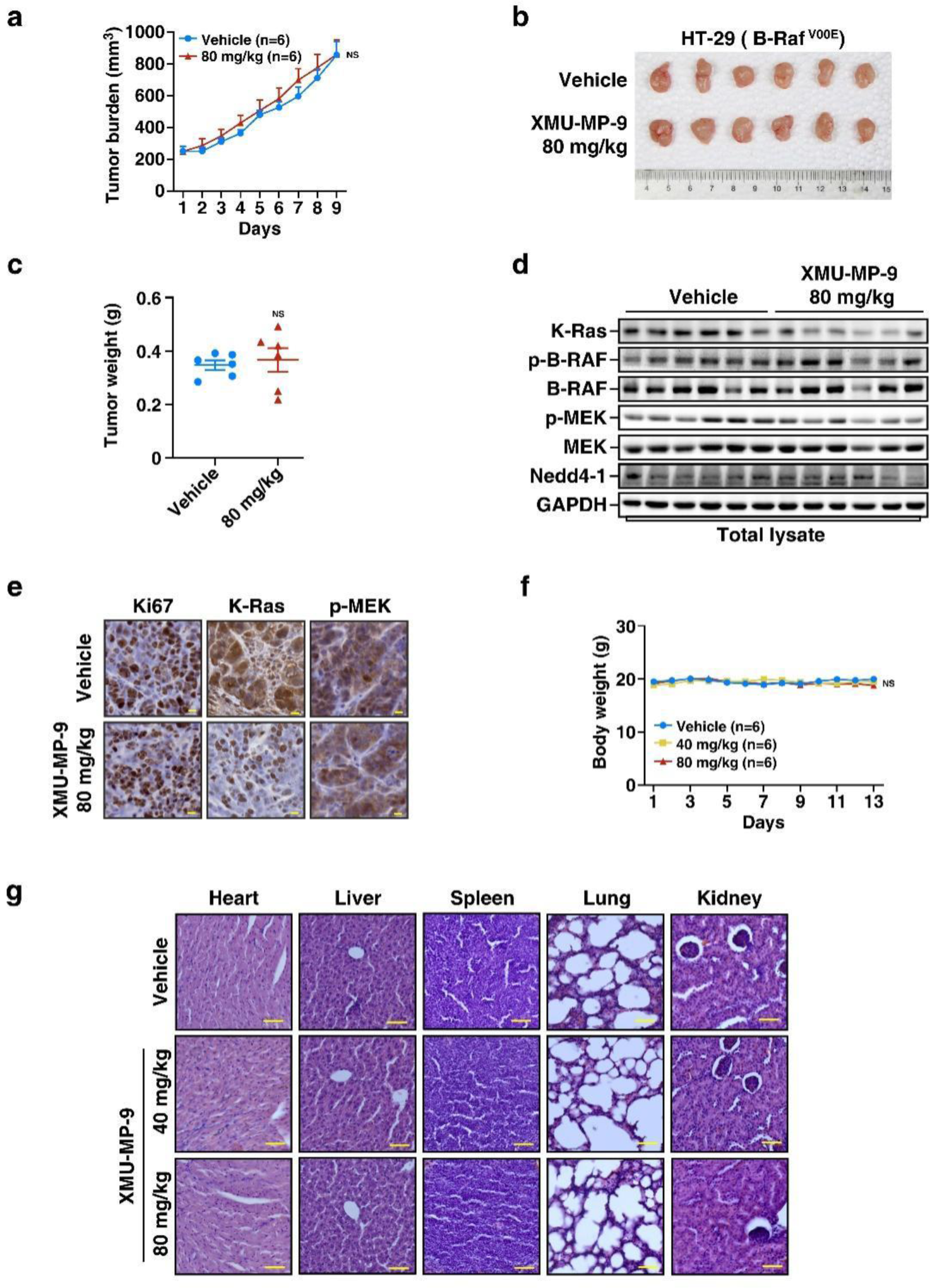
XMU-MP-9 does not inhibit tumor growth of HT-29 cells and has no obvious toxicity to mice. **a-c,** XMU-MP-9 does not inhibit xenograft tumor growth of HT-29 cells in nude mice. HT-29 cells were subcutaneously injected into nude mice to form xenograft tumors. The tumor-bearing mice were given a daily dose of 80 mg/kg of XMU-MP-9 (injected with a dose of 40 mg/kg twice a day via tail vein) or vehicle as control. Tumor volumes were measured and plotted as mean ±SEM (n = 6 animals per group). *P* values were determined by two-way ANOVA followed by Tukey test. NS, not significant (**a**). The tumors were obtained 9 days after drug treatment by sacrificing the mice (**b**), weighted and plotted as mean ± SEM (n = 6 animals per group). *P* values were determined by two-tailed unpaired Student’s t-test. NS, not significant (**c**). **d, e,** Treatment of XMU-MP-9 decreases K-Ras levels but not downstream Raf/MEK signaling in HT-29 cells formed xenograft tumors. Representative tumors were subjected to immunoblotting (**d**) or IHC assay (**e**). Scale bar indicates 10 μm. **f,** XMU-MP-9 does not affect body weights of mice. Body weights of BALB/c mice treated with or without indicated daily dose of XMU-MP-9 were measured and plotted as mean ±SEM (n = 6 animals per group). *P* values were determined by two-way ANOVA followed by Tukey test. NS, not significant. **g,** Organs (heart, liver, spleen, lung, and kidney) of mice treated in (**f**) were collected at the end of the treatment and subjected to H&E staining. Scale bar indicates 50 μm.

## Supplementary Note 1 – Chemical synthesis of XMU-MP-9 and XMU-MP-9-PAL

**Scheme 1.**
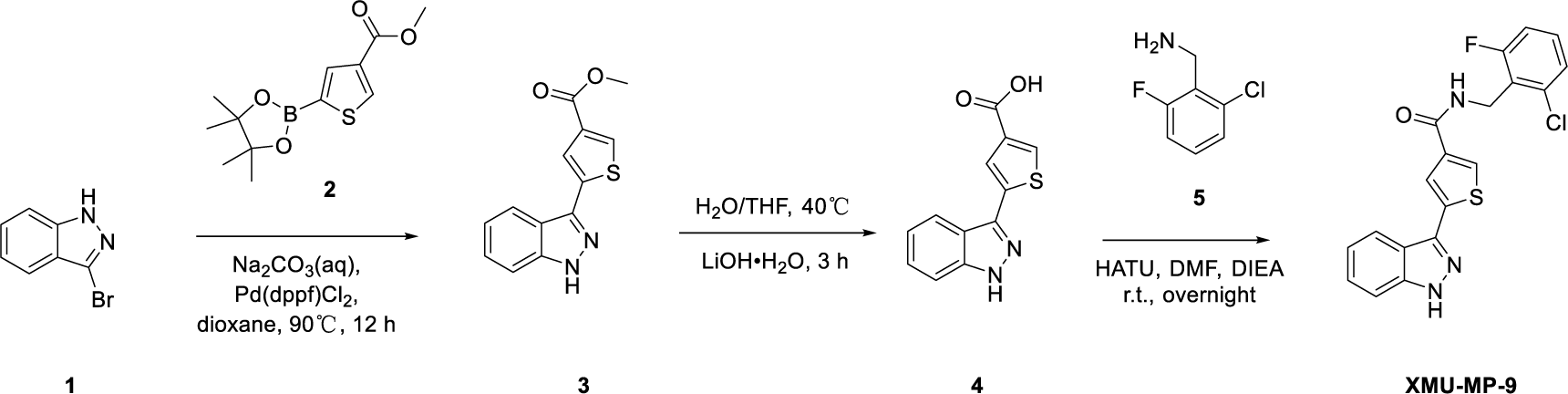
Synthesis of XMU-MP-9.

*methyl 5-(1H-indazol-3-yl) thiophene-3-carboxylate (**compound 3**)*

3-bromo-1H-indazole (**compound 1**, 1.00 g, 5.07 mmol), methyl 5-(4,4,5,5-tetramethyl-1,3,2-dioxaborolan-2-yl) thiophene-3-carboxylate (**compound 2**, 1.63 g, 6.09 mmol) and Pd(dppf)Cl_2_ (0.371 g, 0.508 mmol) were added sequentially to a stirred solution of Na_2_CO_3_ (2.15 g, 20.3 mmol) in 1,4-dioxane (25.0 mL) and H_2_O (25.0 mL). The resulting mixture was heated under argon with vigorous stirring at 90 ℃ for 12 hours until the reaction was completed. Then the reaction was diluted with ethyl acetate (200.0 mL), washed with water (100.0 mL) and brine (50.0 mL), and the organic layer was dried over anhydrous MgSO_4_, filtered and concentrated. The concentrate was purified by flash column chromatography on silica gel with ethyl acetate and hexane (v/v = 15/85) to give **compound 3** (0.26 g, 19.8%). MS (ESI) m/z: 259.2 [M+H]^+^.

*5-(1H-indazol-3-yl) thiophene-3-carboxylic acid (**compound 4**)*

To a stirred solution of **compound 3** (1.10 g, 4.26 mmol) in H_2_O (14.0 mL) and THF (14.0 mL) was added LiOH·H_2_O (1.07 g, 25.55 mmol), then the reaction was heated at 40 ℃ for 3 hours until it was completed. The resulting solution was diluted with water (150.0 mL) and washed with ethyl acetate (50.0 mL). Then the pH value of the aqueous layer was adjusted to 5 by HCl (1.0 M, aq.), and the precipitates was collected and washed with water. The solids were dried by lyophilizer to give **compound 4** (0.97 g, 93.3%), and it was used in next step without further purification. MS (ESI) m/z: 245.2 [M+H]^+^.

*N-(2-chloro-6-fluorobenzyl)-5-(1H-indazol-3-yl) thiophene-3-carboxamide (**XMU-MP-9**)*

A mixture of **compound 4** (50.0 mg, 0.20 mmol), 2-Chloro-6-fluorobenzylamine (**compound 5**, 34.3 mg, 0.21 mmol), HATU (93.4 mg, 0.25 mmol) and DIEA (34.4 mg, 0.27 mmol) in DMF (7.8 mL) was stirred at room temperature overnight until the reaction was completed. The resulting mixture was diluted with ethyl acetate (100.0 mL) and washed with brine (100 mL ×2). The organic layer was dried over anhydrous Na_2_SO_4_, filtered and concentrated under reduced pressure. The residue was purified by flash chromatography on silica gel to give **XMU-MP-9** as a white solid (38.4 mg, 48.7%). ^1^H NMR (600 MHz, DMSO-*d_6_*) δ 13.28 (s, 1H), 8.73 (t, *J* = 5.0 Hz, 1H), 8.15 – 8.10 (m, 3H), 7.59 (d, *J* = 8.4 Hz, 1H), 7.46 – 7.38 (m, 2H), 7.36 (d, *J* = 8.0 Hz, 1H), 7.27 (t, *J* = 7.9 Hz, 2H), 4.61 (d, *J* = 4.8 Hz, 2H). ^13^C NMR (150 MHz, DMSO-*d_6_*) δ 161.7, 161.6 (d, *J* = 249.1 Hz), 141.5, 138.3, 137.7, 136.9, 135.0 (d, *J* = 5.6 Hz), 130.2 (d, *J* = 9.8 Hz), 128.8, 126.8, 125.6 (d, *J* = 3.2 Hz), 123.9 (d, *J* = 17.3 Hz), 122.9, 121.5, 120.4, 119.3, 114.6 (d, *J* = 22.6 Hz), 110.9, 35.0 (d, *J* = 3.8 Hz). MS (ESI) m/z: 386.3 [M+H]^+^. HRMS (ESI, positive mode): calculated for C19H14ClFN3OS [M + H]^+^, 386.0525; found, 386.0516.

**Scheme 2.**
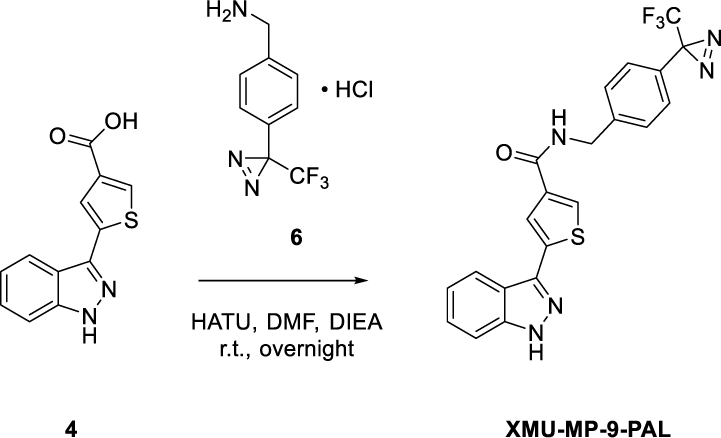
Synthesis of photo-affinity probe XMU-MP-9-PAL.

*5-(1H-indazol-3-yl)-N-(4-(3-(trifluoromethyl)-3H-diazirin-3-yl) benzyl) thiophene-3-carboxamide (**XMU-MP-9-PAL**)*

A mixture of **compound 4** (20.0 mg, 0.08 mmol), (4-(3-(trifluoromethyl)-3H-diazirin-3-yl) phenyl) methanamine hydrochloride (**compound 6**, 21.6 mg, 0.09 mmol), HATU (37.4 mg, 0.10 μμol) and DIEA (24.4 mg, 0.11 mmol) in DMF (2.5 mL) was stirred at room temperature overnight until the reaction was completed. The resulting mixture was diluted with ethyl acetate (100.0 mL) and washed with brine (100 mL × 2). The organic layer was dried over anhydrous Na_2_SO_4_, filtered and concentrated under reduced pressure. The residue was purified by flash chromatography on silica gel to give **XMU-MP-9-PAL** as a white solid (22.2 mg, 61.3%). ^1^H NMR (600 MHz, DMSO-*d_6_*) δ 13.29 (s, 1H), 9.07 (t, *J* = 6.0 Hz, 1H), 8.16 (d, *J* = 8.2 Hz, 1H), 8.15 – 8.13 (m, 2H), 7.62 – 7.58 (m, 1H), 7.49 (d, *J* = 8.3 Hz, 2H), 7.44 (ddd, *J* = 8.1, 6.9, 1.0 Hz, 1H), 7.30 – 7.25 (m, 3H), 4.54 (d, *J* = 5.9 Hz, 2H). ^13^C NMR (150 MHz, DMSO-*d_6_*) δ 161.83, 142.26, 141.41, 138.18, 137.80, 136.98, 128.72, 128.18, 126.63, 126.57, 125.97, 122.54, 121.37, 120.29, 119.18, 110.78, 78.96, 41.94. MS (ESI) m/z: 442.1 [M+H]^+^. HRMS (ESI, positive mode): calculated for C21H15F3N5OS [M+H]^+^, 442.0949; found, 442.0938.

## Supplementary Note 2 – ^1^H and ^13^C spectra of compound XMU-MP-9 and XMU-MP-9-PAL

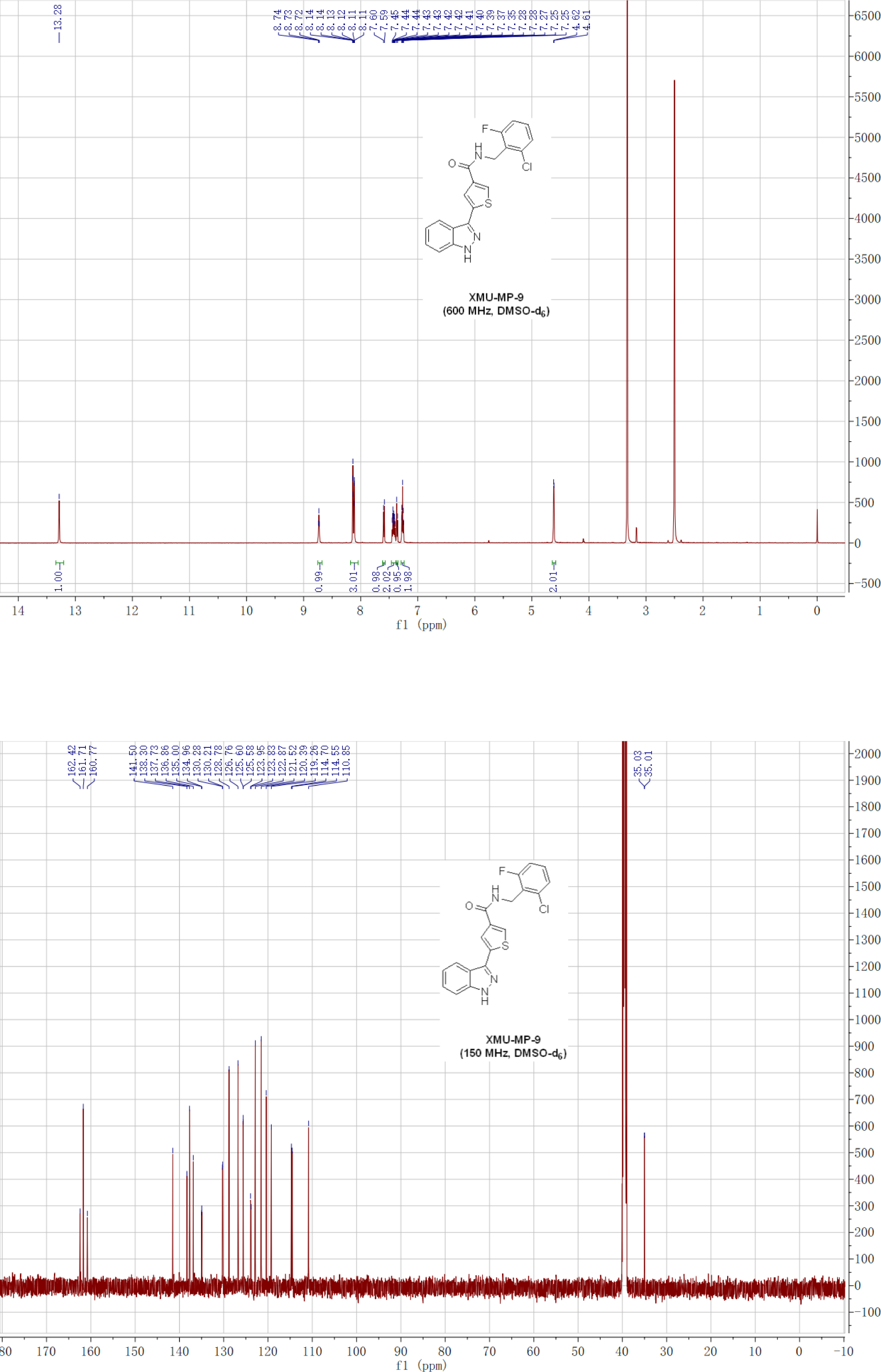
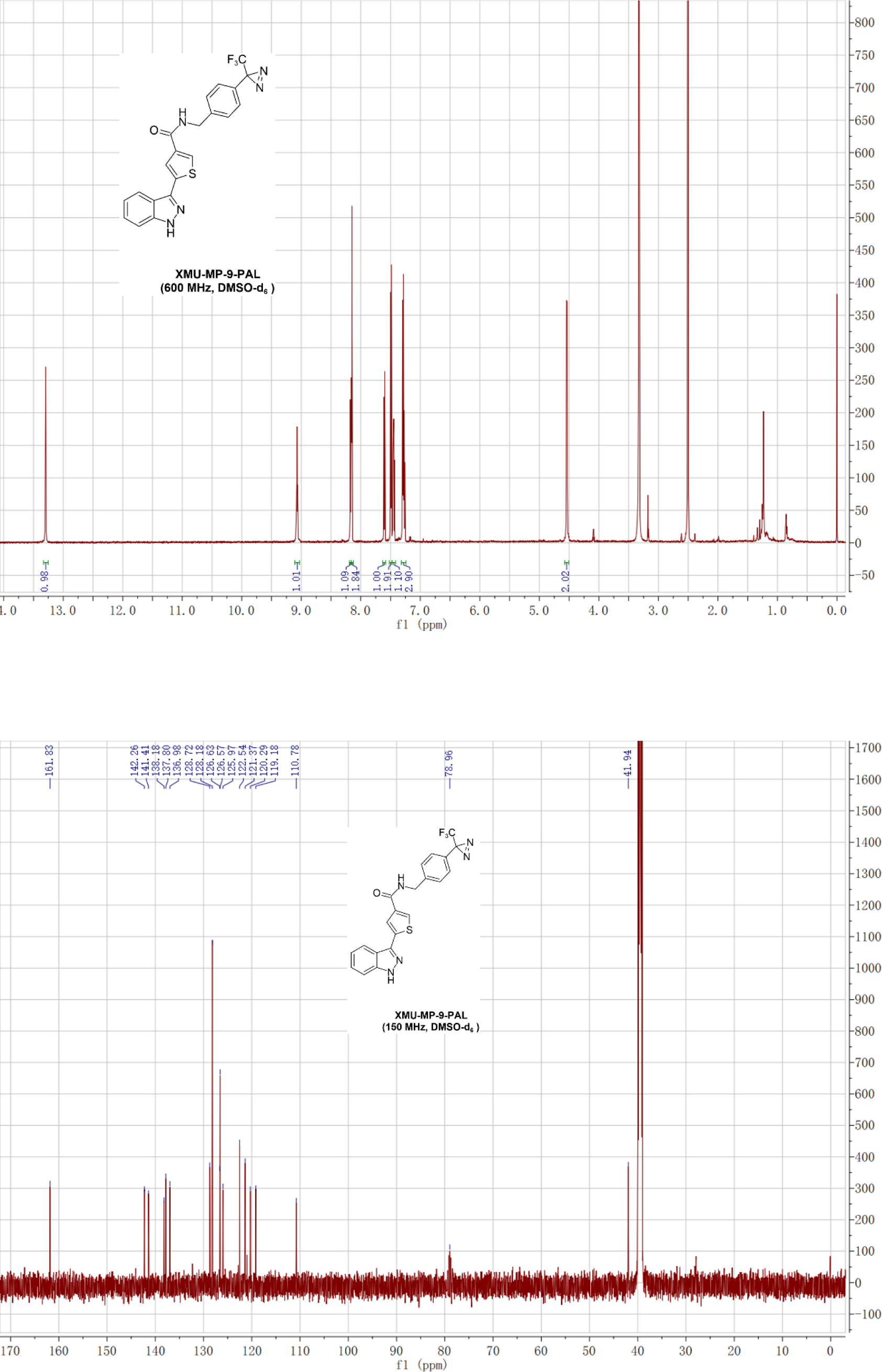

